# FASTiso: Fast Algorithm on Search state Tree for subgraph ISOmorphism in graphs of any size and density

**DOI:** 10.1101/2025.02.28.640915

**Authors:** Wilfried Agbeto, Camille Coti, Vladimir Reinharz

**Affiliations:** Université du Québec à Montréal; École de Technologie Supérieure

**Keywords:** Graphs, graph matching, graph isomorphism, subgraph isomorphism, monomorphism

## Abstract

Subgraph isomorphism is a fundamental combinatorial problem that involves finding one or more occurrences of a pattern graph within a target graph. It arises in a wide range of application domains, including biology, chemistry, social network analysis, and pattern recognition. Although subgraph isomorphism is NP-complete in the general case, many exact algorithms allow it to be solved in practice on many instances. However, the increasing size and structural diversity of graph datasets continue to pose significant challenges in terms of robustness and scalability.

In this article, we propose FASTiso, an exact subgraph isomorphism algorithm that emphasizes a strong consistency between the variable ordering strategy and the pruning rules used during search. This design enables a unified exploitation of structural information throughout the exploration process, leading to improved efficiency and stable performance across heterogeneous graph structures. An extensive experimental evaluation on widely used synthetic and real-world benchmarks shows that FASTiso consistently outperforms reference solvers such as VF3, VF3L, and RI, and achieves competitive performance compared to constraint programming–based approaches (Glasgow, PathLad+), while outperforming them on most datasets. The results further demonstrate that FASTiso remains highly efficient on small instances and scales well to large graphs, while maintaining a lower memory footprint than most evaluated solvers. The peak memory usage is 7.74 GB for FASTiso, 36.19 GB for PathLad+, over 500 GB for Glasgow, 9.62 GB for VF3/VF3L, and 4.31 GB for RI.

FASTiso code is available at https://gitlab.info.uqam.ca/cbe/fastiso as a C++ implementation, a Python module, and an integration within an extended version of NetworkX. The implementations support simple graphs and multigraphs, directed or undirected, with labels on nodes, edges, or both.

## I. Introduction

Graphs are present in many fields, such as biology, chemistry, bioinformatics, social networks, databases, and pattern recognition. They are used to model data for better analysis and description, which often involves searching for one or all occurrences of a smaller graph (pattern graph) within a larger graph (target graph). This problem is known in the literature as the subgraph isomorphism problem. For example, in biology, molecular structures such as RNA can be represented as graphs, where nodes represent nucleotides and edges represent interactions between them. RNA exhibits a highly modular structure with essential motifs that influence the three-dimensional configuration of the molecule and, consequently, define its role in the cell. Understanding these modules primarily requires identifying subgraphs with specific topologies, thus involving the use of subgraph isomorphism algorithms [10], [45], [55], [2]. In chemistry, subgraph isomorphism is used to identify similar chemical structures within large molecular databases, which can be useful for drug design and the prediction of chemical reactivity. Some tools [25], [62] used in bioinformatics or for code analysis incorporate subgraph isomorphism solvers, so the performance of these tools depends on the efficiency of the solver used. The subgraph isomorphism is also widely used in Pattern Recognition [60], semantic search [65], RDF query processing [30], symbol recognition [39], and community detection [13].

Subgraph isomorphism is an NP-complete problem [21]. Due to this complexity, even though subgraph isomorphism has numerous and varied applications, finding efficient solutions remains challenging. Most exact algorithms rely on two key approaches to improve efficiency [9]. First, a variable ordering strategy is used to determine the order in which nodes of the pattern graph are matched during the search. This choice has a major impact on the size of the search tree, yet no universally effective ordering strategy has been proposed so far. Second, pruning or inference rules are employed to detect early that a partial mapping cannot be extended into a valid solution. While powerful pruning techniques can dramatically reduce the search space, they often come at a high computational cost and may be effective only for specific graph structures. As a result, no single algorithm consistently outperforms all others across heterogeneous datasets.

Additional factors, such as graph size and density, further influence the practical complexity of subgraph isomorphism. Several approaches [18], [17], [5] have been specifically designed for large graphs (with more than 1,000 nodes) or dense graphs (with density greater than 0.1). Nevertheless, the rapid growth of modern graph datasets, both in size and structural complexity, calls for continuous improvements to existing methods, particularly with respect to robustness and scalability.

### Our contributions

Performance instability across existing subgraph isomorphism algorithms may arises from a suboptimal synergy between variable ordering strategies and pruning rules. For instance, VF3 employs sophisticated pruning mechanisms based on a partitioning of the node neighborhood, yet this information is not fully exploited to guide node exploration. Conversely, RI leverages a similar neighbor partitioning to determine the exploration order but does not explicitly use this information to strengthen pruning.

As a consequence, the potential synergy between variable ordering and pruning remains largely underexploited. In practice, although both aim to reduce the search space, the variable ordering has a direct influence on the effectiveness of the pruning rules applied during the search. These observations suggest that robustness across heterogeneous graph datasets can be improved by tightly integrating variable ordering and pruning strategies, designing them jointly to exploit the same structural information for greater combined effectiveness.

Motivated by the need to improve the performance of exact subgraph isomorphism algorithms on heterogeneous datasets in terms of robustness and scalability, we propose FASTiso, a new exact algorithm using the VF3 algorithm as a reference. FASTiso introduces two new pruning rules that are both effective and computationally inexpensive, alongside a variable ordering strategy explicitly aligned with the pruning rules used to reduce the search space. This ensures that the same structural information guides both the exploration order and the elimination of unpromising search branches. We integrate these contributions into an exact solver derived from VF3, significantly improving robustness and scalability across diverse datasets.

FASTiso was compared to VF3 and other state-of-the-art solvers (VF3L [15], RI [11], Glasgow [42], and PathLad+ [61]) on widely used datasets, including both synthetic and real graphs. We also evaluated the scalability of FASTiso on very large graphs containing up to 23 million nodes. The results show that FASTiso significantly reduces matching time compared to VF3 and the other solvers while maintaining high generality and scalability. Further experimental analyses confirm that the proposed strategies strongly contribute to FASTiso’s superior performance across diverse datasets.

The paper is structured as follows. Section II reviews previous work on subgraph isomorphism. In Section III, we first formalize the problem and then present the FASTiso algorithm. Finally, Section IV presents the experimental results.

## II. Related works

Several exact algorithms have been developed over time to solve the subgraph isomorphism problem. Most of these approaches can be classified into three main categories: tree search, constraint programming, and graph indexing [18], [63].

Tree search methods define the subgraph isomorphism problem as a state space exploration problem, typically solved using a depth-first search technique with backtracking. Each state represents a partial solution, meaning a partial mapping between the nodes of the pattern graph and the target graph. The goal is to find valid solutions while minimizing the exploration of invalid states. These algorithms begin with an empty initial state and progressively add pairs of nodes that satisfy the subgraph isomorphism constraints until a feasible solution is found.

Pruning rules play a crucial role in the efficiency of a tree search-based algorithm. Some algorithms, such as VF2, VF3, incorporate lookahead functions after verifying that the current state satisfies the conditions of subgraph isomorphism. These functions assess whether the current state cannot lead to a valid solution, thereby reducing the number of explored states by avoiding partial solutions that are bound to fail.

Generally, these functions rely on the classification of nodes to detect inconsistencies. The analysis is usually performed locally, searching for inconsistencies among the neighbors of the node pairs being matched. However, the computational cost of a lookahead function is proportional to its ability to identify inconsistencies. When the graphs contain inconsistencies that the function cannot detect, the additional computational cost becomes unnecessary overhead. Therefore, a trade-off is necessary between the cost of the lookahead function and its effectiveness in reducing the search space. For this reason, some algorithms, such as RI, do not use lookahead functions or use very lightweight ones and instead prioritize an optimal variable ordering.

It is important to have a variable ordering strategy to guide the search and eliminate unfruitful states in the early stages of exploration. This relies on consistency between the rules used to determine this order and those applied to prune the search space. The order in which the nodes of the target graph are selected to be matched to the nodes of the pattern graph, called the value ordering, can also be defined.

Ullmann’s algorithm [58] was the first to adopt this approach, but it took more than 20 years before a truly efficient algorithm, namely VF2 [22], was proposed. Several improvements to VF2 have since been developed, such as VF2 Plus [19], VF2++ [31], and VF3 [18], [17]. Other notable algorithms include RI/RI-DS [11], RI+ [4], VF3L [15], and L2G [3]. VF2 Plus and RI/RI-DS are particularly effective for biological data, but VF2 Plus is faster on large graphs. VF3 is better suited for large and dense graphs, and VF3L is an enhancement of VF3 that omits certain pruning rules that are less effective on large and dense graphs.

Subgraph isomorphism can also be solved using Constraint Satisfaction Programming (CSP), which generally involves assigning a value from the domain of pre-defined variables (the possible values that a variable can take) while respecting a set of constraints. The resolution of this problem often relies on inference techniques combined with search strategies using backtracking. The constraints limit the valid combinations of values that subsets of variables can take. Thus, when a value is assigned to a variable, inference consists of propagating the constraints involving that variable to reduce the domains of other variables by removing values that are invalid or inconsistent with the propagated constraints.

In the case of subgraph isomorphism, the nodes of the pattern graph are treated as variables, while the nodes of the target graph form the domain. The constraints are those imposed by the conditions of subgraph isomorphism. Initially, before the search phase, a compatibility domain is calculated for all nodes of the pattern graph using filtering techniques based on the degree and label of the nodes. More advanced filtering techniques can also be used [68], [40].

During the search, inference techniques are employed to propagate constraints and reduce the domains of the variables. These methods generally use more memory than tree search methods because they store compatibility domains and incompatibilities arising from constraint propagation to avoid re-evaluating them. Inference can be either global or local, but in both cases, unlike methods based on tree search, it enables the detection of inconsistencies involving nodes that are not neighbors of the pair of nodes currently being matched. However, this comes at the cost of increased computational effort. In the presence of graphs with symmetrical or specific structures, where constraints can significantly reduce the search space, constraint programming-based methods can be more efficient, whereas tree search-based methods may suffer from weaker prediction capabilities. However, there exist instances of the subgraph isomorphism problem that remain particularly challenging for both approaches [53], [41].

The first constraint programming-based algorithm to solve the subgraph isomorphism problem was proposed by McGregor [43]. Later algorithms such as nRF+ [36], ILF [68], LAD [51], PathLad [33], PathLad+ [61], FocusSearch [59], SND [6], and Glasgow [42] followed.

The graph indexing approach, on the other hand, is used to efficiently search large graph databases for those that contain a given pattern. Generally, indexing methods build an index representing the structural and semantic characteristics of graphs. The index is used to filter target graphs by quickly eliminating those that cannot contain the searched pattern. Then, subgraph isomorphism constraints are verified on the remaining graphs. Algorithms that use this method include GraphQL [29], QuickSI [49], SPath [70], GADDI [69], CFL-Match [8] TurboIso [28], CECI [7], DP-iso [26], and CFQL [56].

Other works have addressed the non-induced subgraph isomorphism problem to reduce redundant exploration caused by symmetries in graphs. These approaches exploit symmetries to compress equivalence classes [27], [46], [44], [32], [67]. Portfolio-based approaches have also been proposed to dynamically select the best algorithm from a set of solvers for a given instance [33], [57].

Parallel computing has been widely explored to accelerate subgraph isomorphism, first on multi-core CPUs [63] and more recently on GPUs [24] or hybrid CPU–GPU frameworks [20]. However, these approaches often face scalability issues due to the irregular search space, which leads to severe load imbalance and costly communication for dynamic workload redistribution. The inherent complexity of parallel approaches highlights the continued importance of advancing sequential exact algorithms. Improvements in pruning rules and ordering strategies directly reduce the search space explored by parallel methods, thereby mitigating load imbalance. Moreover, robust sequential solvers provide essential baselines for a fair evaluation of the speedup and scalability of parallel algorithms.

Given the NP-completeness of the problem, approximate or inexact algorithms have also been proposed for scenarios where finding an optimal solution is not strictly required, and a sufficiently close solution is acceptable. These approaches include methods based on probabilistic relaxation [64], [66] as well as learning-based approaches using graph neural networks (GNNs) [48], [34], [38], [35], [50].

Beyond exact one-to-one mapping, some studies have explored more flexible subgraph isomorphism formulations [37], [12]. For example, Lin et al. [37] proposed a generalized subgraph query model in which query edges may be mapped to paths in the data graph under distance constraints. Their approach is designed for large-scale database environments and relies on indexing and a cost model to compute the variable ordering efficiently.

## III. Method

### A. Background

A graph *G* is defined by a set *V* of nodes and a set or multiset *E* ⊆*V* × *V* of edges connecting these nodes. The order of a graph, denoted |*V* |, refers to the number of nodes, while its size, denoted |*E*|, corresponds to the number of edges. A graph can be directed or undirected. In an undirected graph, an edge {*u, v*} represents a bidirectional relationship between nodes *u* and *v*. In a directed graph, an edge (arc) (*u, v*) represents a unidirectional relationship that goes from *u* to *v*. In this case, *u* is called the predecessor of *v*, and *v* is called the successor of *u*. A graph can also be labeled if labels are assigned to its nodes or edges. The labeling functions *λ* : *V* → *A*_*V*_ and *β* : *E* → *A*_*E*_, respectively, associate to each node and edge the corresponding labels.

For the remainder of this article, the focus will be on directed graphs. However, the proposed approach remains applicable to simple undirected graphs, as well as to both directed and undirected multigraphs.

The out-neighborhood, and in-neighborhood of a node *u* are defined as

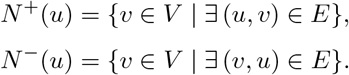

The (total) neighborhood of *u* is *N*(*u*) = *N* ^+^(*u*) ∪ *N* ^*−*^(*u*). Neighborhoods are defined as sets of distinct nodes and therefore ignore edge multiplicities. The out-degree of *u*, denoted *d*^+^(*u*), is defined as *d*^+^(*u*) = | {(*u, v*) ∈ *E | v* ∈ *V*}|, that is, the number of outgoing edges incident to *u*, counting multiplicities. Similarly, the in-degree of *u*, denoted deg^*−*^(*u*), is defined as *d*^*−*^(*u*) = |{(*v, u*) ∈ *E* | *v* ∈*V* }|. The total degree of *u* is then given by *d*(*u*) = deg^+^(*u*) + deg^*−*^(*u*).

Let *G* = (*V, E*) be a pattern graph and *G*′ = (*V* ′, *E*′) be a target graph. The subgraph isomorphism problem consists of finding an injective function *M* : *V*→*V* ′ that maps each node of *G* to a unique node of *G*′ (condition 1-2 of the subgraph isomorphism). Moreover, for every pair of nodes *u, v* in *G*, and for every pair of corresponding nodes *u*′ = *M*(*u*), *v*′ = *M*(*v*) in *G*′, if an edge (*u, v*) exists in *G*, then there must be a corresponding edge (*u*′, *v*′) in *G*′; if an edge (*u, v*) does not exist in *G*, then there is no corresponding edge (*u*′, *v*′) in *G*′ (condition 3-4 of the subgraph isomorphism). If the nodes or edges are labeled, then the labels must be preserved (condition 5-6 of the subgraph isomorphism). Thus, we have the following conditions:

1. ∀*u* ∈ *V*, ∃*u*′ ∈ *V* ′ : (*u, u*′) ∈ *M*
2. ∀*u, v* ∈ *V, u* ≠ *v* ⇒ *u*′ ≠*v*′
3. ∀(*u, v*) ∈ *E*, ∃(*u*′, *v*′) ∈ *E*′
4. ∀*u, v* ∈ *V*, (*u*′, *v*′) ∈ *E*′ ⇒ (*u, v*) ∈ *E*
5. ∀*u* ∈ *V*, 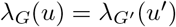
6. ∀(*u, v*) ∈ *E*, 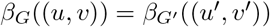

It is possible that edges in *G*′ do not have a corresponding match in *G*; in this case, the subgraph isomorphism is referred to as a monomorphism (non-induced subgraph isomorphism) [21] and can be defined by removing condition 4 above.

### B. The foundation of FASTiso

FASTiso, like its predecessor VF3, is based on a tree search method that reformulates the subgraph isomorphism problem as an exploration of a state space. The set of possible states can be represented by a search state-space tree (SST) (see supplementary file, Figure F). Each node of this tree represents a possible match between a node *u* of *G* and a node *u*′ = *M*(*u*) of *G*′. The path from the root to a given node in this tree represents a partial mapping 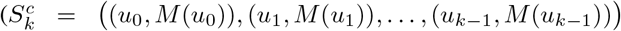, with *k <* |*V* |) between *G* and *G*′. We define 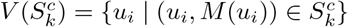 and 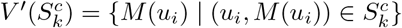 as respectively the set of nodes of *G* and *G*′ that are part of the partial state 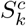. The partial state 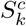 induces two subgraphs of *G* and *G*′, defined respectively as

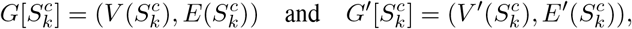

where 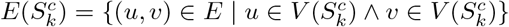, and 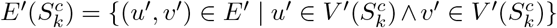. Using the terminology of VF3, a partial mapping (partial state) 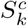 is said to be a consistent state if it satisfies all the conditions of the subgraph isomorphism problem, that is, if the induced subgraphs 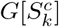 and 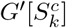 satisfy consditions 1–6 and are therefore isomorphic. A consistent state that includes all nodes of *G* (i.e., *k* = |*V* |) is called a goal state and represents a complete solution to the subgraph isomorphism problem. Conversely, a consistent state that cannot be extended to any goal state is called a dead state.

The Algorithm starts from an initial state 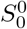 (the empty state *∅*) and traverses the consistent states by progressively adding ordered pairs (*u*_*k*_, *M*(*u*_*k*_)) such that 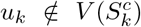 and 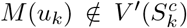 until a solution or feasible solutions (goal states) are found. One way to improve the algorithm’s performance is to reduce the number of states visited by identifying dead states as early as possible. This can be achieved by selecting a good variable ordering and using pruning rules.

Algorithm 1 presents the foundation of FASTiso. The first step consists of computing an initial domain assignment for each node of the pattern graph, considering their label, degree, and the degree of their neighbors. We define a compatibility map *C* between the nodes *u*_*i*_ of the pattern graph and the nodes 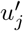 of the target graph, such that:

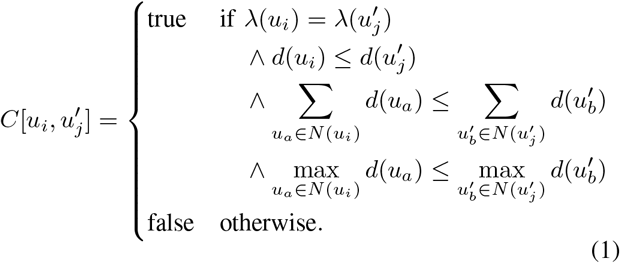

A node 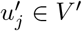 is said to be compatible with a node *u*_*i*_ ∈ *V* if *u*_*i*_ and 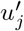 have the same label, the degree of *u*_*i*_ is less than or equal to the degree of 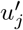, the sum of the degrees of the neighbors of *u*_*i*_ is less than or equal to that of 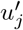, and the maximum degree among the neighbors of *u*_*i*_ is less than or equal to that of 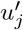.

We also define a list d size that contains the domain size for each node *u*_*i*_. Next, we compute the variable ordering (function computeOrdering, more details in Section III-C), and finally, we explore the SST using a depth-first search with backtracking to find the goal states (function match, more details in Section III-D).

#### Algorithm 1 FASTiso

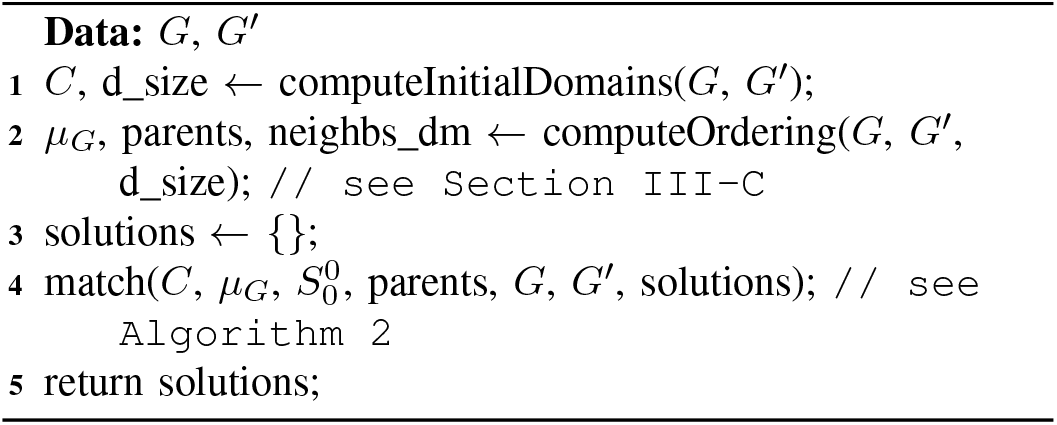

The improvement introduced by FASTiso compared to other tree search methods primarily lies in the variable ordering strategy and pruning rules. Regarding variable ordering, we introduced new rules that are strongly aligned with the pruning rules used to reduce the search space, allowing better node discrimination and a more effective prioritization of pattern vertices during the search.

For pruning rules, we introduced two new approaches to enhance the algorithm’s efficiency.

(1) Verification of conditions 3 and 4 of the subgraph isomorphism can be computationally expensive (see Section III-D). To address this, we incorporated a low-cost filtering test, which avoids systematically evaluating these conditions for every node in the SST. (2) We proposed a lookahead function that enables faster detection of dead states.

### C. Variable Ordering Strategy

An overview of the different strategies for calculating the variable ordering can be found here [9]. Generally, there are two types of variable ordering: static variable ordering, where the order in which the pattern graph nodes are mapped is determined at the beginning and remains fixed during the SST visit, and dynamic variable ordering, which involves deciding, as the SST visit progresses, which node of the pattern graph to map. These two types of variable ordering can be computed based on information about the structure of the pattern graph and the target graph.

The variable ordering can be seen as a permutation of the nodes in the pattern graph. The goal is to find the permutation that allows to quickly determine the non-consistent or dead states. To achieve the aforementioned goal, the logic is to prioritize, i.e., place first in the variable ordering, the nodes of the pattern graph with the most verifiable constraints during the SST visit. This helps to quickly reduce the search space by eliminating invalid matches early on.

Let *µ*_*G*_ be a variable ordering. Our strategy for computing *µ*_*G*_ is based on a combination of the approaches used by VF3 and RI, which we consider complementary while introducing notable improvements. During our experiments with VF3, we observed that the selection of nodes to determine the variable ordering was often done randomly, due to the lack of additional rules to better discriminate the nodes. This led to poor performance on certain datasets. To address this drawback, we decided to combine the methods of VF3 and RI to achieve a more relevant and efficient variable ordering.

It is well-established that the selection of the first node to include in a variable ordering significantly impacts the quality of the resulting order [9]. Our strategy for determining the first node is based on two criteria: if the graph is unlabeled, we select the node with the highest degree; otherwise, we choose the node with the smallest compatibility domain.

The next nodes to be added to the partial variable ordering *µ*_*G*_(*k*) are selected based on their neighbors and their compatibility domain. Let *u*_*i*_ be a candidate node to be added to *µ*_*G*_(*k*), its neighbors *N*(*u*_*i*_) can be classified into three groups:

- *N*_1_(*u*_*i*_) = {*u*_*j*_ : *u*_*j*_ ∈*N*(*u*_*i*_) and *u*_*j*_ ∈*µ*_*G*_(*k*) }, the set of neighbors that are present in the partial ordering.
- *N*_2_(*u*_*i*_) = {*u*_*j*_ : *u*_*j*_ ∈*N*(*u*_*i*_) and *u*_*j*_ ∈*N*(*µ*_*G*_(*k*)) }, the set of neighbors that are not in the partial ordering but have at least one neighbor that is.
- *N*_3_(*u*_*i*_) = { *u*_*j*_ : *u*_*j*_ ∈*N*(*u*_*i*_) and *u*_*j*_ ∉*N*_1_(*u*_*i*_) and *u*_*j*_ ∉*N*_2_(*u*_*i*_) }, the set of neighbors that do not belong to *N*_1_(*u*_*i*_) or *N*_2_(*u*_*i*_).

Let *u*_*i*_, *u*_*j*_ be two candidate nodes to be added to *µ*_*G*_(*k*) and *N*_edges_(*u*_*i*_, *S*) the total number of incoming and outgoing edges between a node *u*_*i*_ and the node in the set *S*:

A. *u*_*i*_ is selected if:

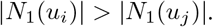
B. In case of equality, *u*_*i*_ is selected if:

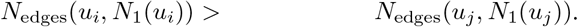
C. in case of equality, *u*_*i*_ is selected if:

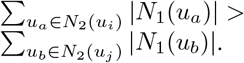
D. in case of equality, *u*_*i*_ is selected if:

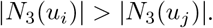
E. in case of equality, *u*_*i*_ is selected if:
  - *d*_size_[*u*_*i*_] *> d*_size_[*u*_*j*_], if *G* is unlabeled,
  - *d*_size_[*u*_*i*_] *< d*_size_[*u*_*j*_], if *G* is labeled.
F. in case of equality, *u*_*i*_ is selected if:

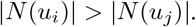

FASTiso Choose by priority order: (1) The node with the most mapped neighbors, or the node whose neighbors have the most mapped neighbors. (2) The node with the smallest domain if the graph is labeled, and if the graph is unlabeled, the node with the largest domain. (3) The node with the highest degree. Figure A (see supplementary file) illustrates an example of the variable ordering computation for the graph on the left, using our strategy.

For each node *u*_*i*_ added to *µ*_*G*_, we store its parent (its first neighbor that was added to *µ*_*G*_) as well as 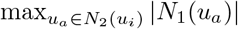. Let parents be the list of parents and neighbs _dm be the list of 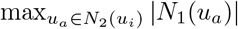 for each node *u*_*i*_. We will see in the next section the usefulness of these lists.

### D. The State-Space Tree (SST) Exploration

Starting from the initial state 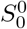 (the empty state ∅), each step involves selecting, at each level *k* of the SST, the candidate nodes 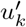 in the target graph to match with the node *u*_*k*_ (the *k*-th node in the variable ordering *µ*_G_).

The candidate nodes are chosen from among the neighbors of the node corresponding to the parent of *u*_*k*_ that are within the compatibility domain of *u*_*k*_ (i.e., 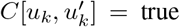, see Equation 1). As shown in Algorithm 2, Line 7-8, when *u*_*k*_ does not have a parent, the set of candidates is the set of all nodes in the target graph. This situation occurs either when *u*_*k*_ is the first node in the variable ordering or when the graphs have multiple connected components.

Before adding the pair 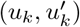 to the state 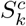 (partial mapping), FASTiso uses feasibility rules (function isFeasible Algorithm 2, Line 13) to check whether this addition will result in a new state that satisfies the constraints of subgraph isomorphism and predict whether it can not lead to a goal state.

The feasibility rules used by FASTiso rely on feasibility sets (defined below) as well as 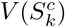 and 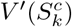, which respectively represent the set of nodes in the pattern graph and the target graph that are already mapped. For a node *u*, the feasibility sets are the three sets of classified neighbors, which are analogous to those used for computing the variable ordering (Section III-C, as we have said, there must be some consistency between the rules used for computing the variable ordering and those used for pruning the search space):

- 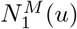: the set of neighbors of *u* that are already mapped,
- 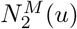: the set of neighbors of *u* that are not yet mapped but have neighbors that are already mapped,
- 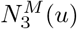: the set of neighbors of *u* that belong neither to 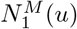 nor 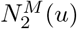.

#### Algorithm 2: match

traverses the search state-space tree using a depth-first search with a backtracking mechanism.

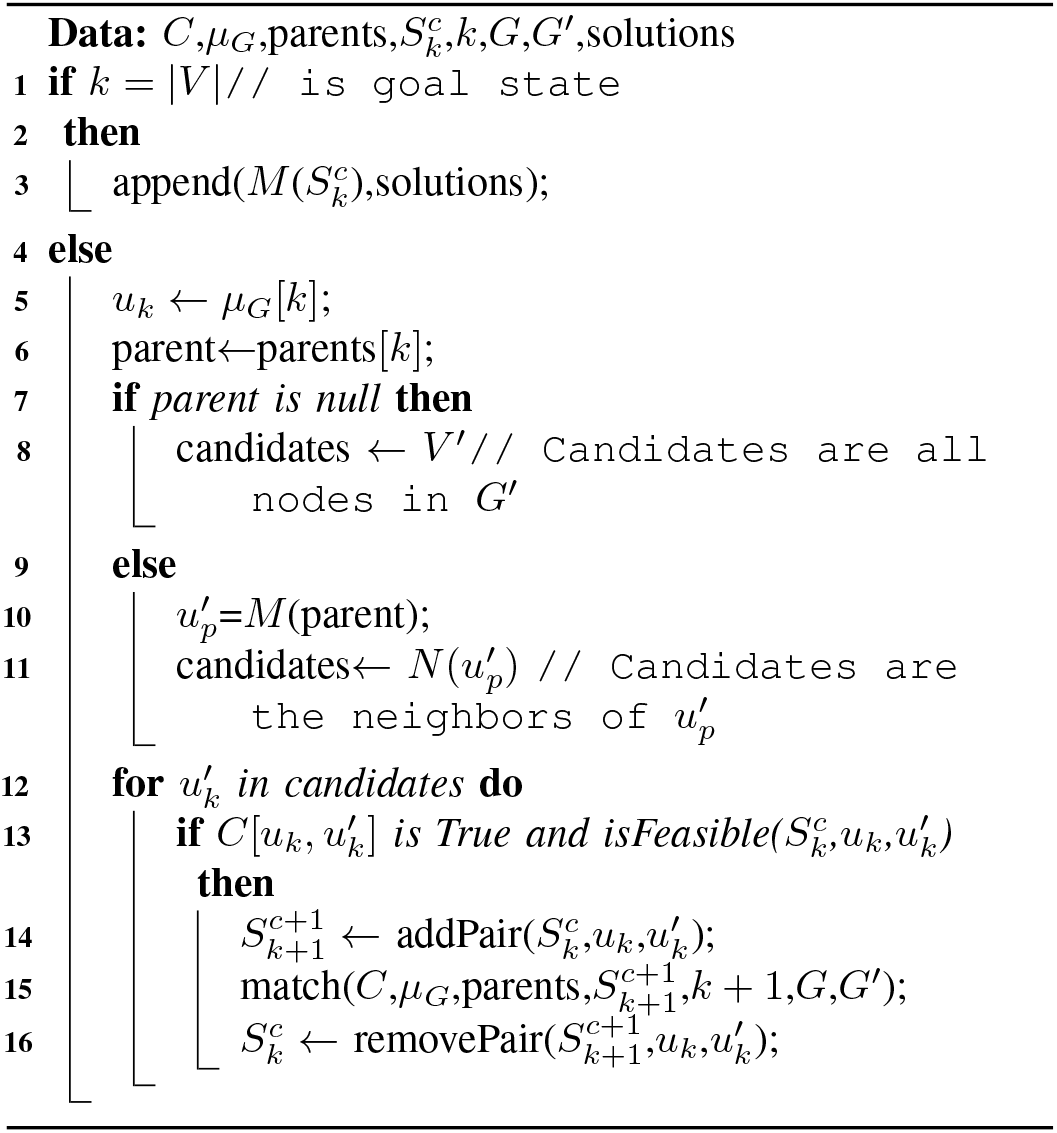

VF3 also uses these feasibility sets, but among the neighbors of *u*, it distinguishes between predecessors and successors.

The functionm isFeasible, which verifies feasibility rules, is formally defined as follows:

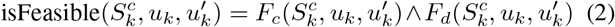

#### Component F_c_

Verify consistent states.

For every pair 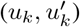 to be added to 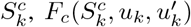 checks the following conditions:

1. 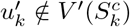, verify condition 2 of the subgraph isomorphism.
2. 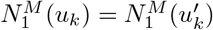,
3. 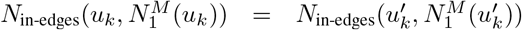 and 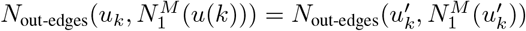
4. For every {*u*_*k*_, *u*_*a*_} ∈ *E* where 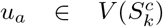, there exists 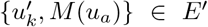 and 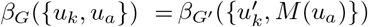, verify conditions 3, 4, and 6 of the subgraph isomorphism.

The conditions 1 and 4 of 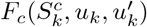 are sufficient to verify the subgraph isomorphism requirements. It is worth noting that condition 4 of subgraph isomorphism is not explicitly verified due to conditions 2 and 3 of 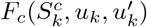, which ensure that the number of neighbors, incoming edges, and outgoing edges of *u*_*k*_ in 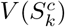 are equal to those of 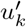 in 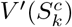. VF3 does not incorporate condition 2, so it explicitly evaluates the conditions 3 and 4 of the subgraph isomorphism.

Checking the presence of an edge can be computationally expensive (especially for dense graphs or during frequent evaluations). For a node *u*, this check requires Θ(*N*(*u*)) using an adjacency list. If the data is sorted, it can be done in Θ(log(*N*(*u*))). Of course, if an adjacency matrix is used, the check can be performed in Θ(1). However, like VF3 and RI, FASTiso uses an adjacency list for memory efficiency, the verification of condition 4 of 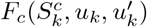 will therefore be done in 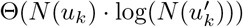.

Conditions 2 and 3 of 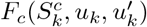 serve as filtering tests to avoid evaluating condition 4 of 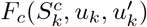 at each node of the SST, which can lead to substantial computational costs.

Condition 5 of subgraph isomorphism is not verified in 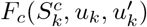 because it is already verified during the computation of compatibility domains (see Section III-B).

To strengthen the filtering tests (conditions 2 and 3), additional information about the identity of the nodes is introduced to minimize the need to evaluate condition 4 as much as possible (see supplementary file, Figure B).

Suppose the nodes are also assigned an integer identifier corresponding to their index, e.g., ID(*u*_1_) = 1. We define a new condition:

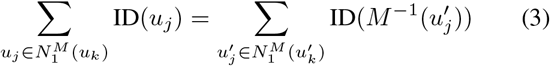

which verifies for every new pair 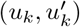 whether the sum of the identifiers of *u*_*k*_’s neighbors already mapped equals the sum of the identifiers of the correspondents of the already mapped neighbors of 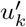.

The implementation details of the component *F*_*c*_ are described in the supplementary material (see Section C).

#### Component 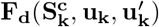

Verify dead states. 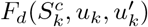 uses lookahead functions that are necessary but not sufficient conditions. When these conditions are false, they guarantee that adding the pair 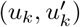 to 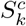 will not lead to a goal state.

For every pair 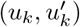 to be added to 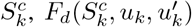 checks the following conditions:

1. 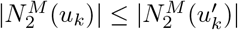
2. 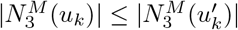

The function *F*_*d*_ can be strengthened by partitioning the nodes to detect more inconsistencies (see supplementary file, Figure C). This methodology is already used by VF3, where nodes are partitioned based on their labels. As a result, this partitioning approach can only be applied to labeled graphs. In our case, we use a classification function that, for a node *u*, partitions 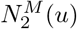 based on the number of their neighbors that are mapped.

For a node *u*, the sets of its classified neighbors are defined as: 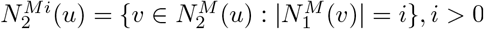

For example, the set 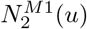 represents all the neighbors of *u* that are not yet mapped but have exactly one mapped neighbor.

For a node *u*, there can be at most 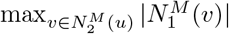 classes. This value is recorded in the list neighbs_dm (see Section III-C) for the nodes of the pattern graph. For every pair 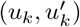 to be added to 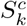, we introduce a new condition to *F*_*d*_ based on classified neighbor sets. The condition is:

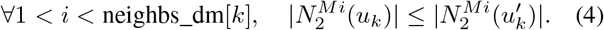

If 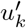 has *j* classes and *u*_*k*_ has *i* classes with *j > i*, there is no need to verify classes *i*+1, *i*+2, …, *j*, as the cardinality of these classes for *u*_*k*_ is zero. This new condition is consistent with the condition (C) of our variable ordering.

The implementation details of the component *F*_*d*_ are described in the supplementary material (see Section D). We have also proposed versions for graph isomorphism and monomorphism (non-induced subgraph isomorphism) for isFeasible 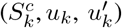 (see supplementary file, Section E).

## IV. Results

To evaluate the performance of FASTiso, we compared it with VF3, VF3L, RI, Glasgow, and PathLad. We selected these solvers because they currently represent leading methods for solving the subgraph isomorphism problem [53], [14]. VF3, VF3L, and RI are also based on tree search, while Glasgow and PathLad are based on constraint programming. All of these solvers, like FASTiso, are implemented in C++. Most graph indexing approaches focus on the non-induced subgraph isomorphism problem and are therefore excluded from our experiments. Similarly, approaches based on graph neural networks (GNNs) are not considered, since they do not guarantee finding exact solutions.

### A. Datasets

Subgraph isomorphism is an NP-complete problem whose difficulty depends less on the size of the graphs than on their structural properties. For this reason, we use in our experiments a diverse collection of datasets, including both real-world and synthetic data. Our datasets are divided into three benchmarks.

#### 1) Classic benchmark [53]

The first benchmark is composed of relatively small graphs containing up to 10,000 nodes (see supplementary file, Figure E). These datasets include both sparse and dense graphs derived from real-world and synthetic sources.

- **MIVIA LDG [15]:** MIVIA Large and Dense Graphs contains large, dense random Erdős–Rényi graphs, both labeled (uniform and non-uniform) and unlabeled, with densities of 0.2, 0.3, and 0.4. Target graphs contain 300–10,000 nodes, and motifs contain 200–2,000 nodes.
- **Scalefree [52]:** A dataset of scale-free networks generated using a power-law degree distribution. Target graphs contain 200–1,000 nodes, while motif graphs represent 90% of the corresponding target graph size. Some motif–target pairs in the dataset have no solution.
- **Phase [41]:** This dataset consists of 200 randomly generated graph pairs near the satisfiable–unsatisfiable phase transition. Motif graphs have 30 nodes and target graphs 150 nodes, with all pairs being unsatisfiable. For our experiments, we used the first 50 pairs.
- **SI dataset (RAND, BVG, M4D, M4DR) [52]:** This dataset includes bounded-valence graphs, random graphs, and 4D meshes, with target graphs containing 200–1,296 nodes and motifs 20–60% of the target size, sourced from the MIVIA dataset [1].
- **Images-PR15 [52], [54]:** contains graphs generated from segmented images, with a target graph of 4,838 nodes and 24 motif graphs ranging from 4 to 170 nodes.
- **Images-CVIU11 [52], [23]:** contains graphs generated from segmented images, with motifs of 15–151 nodes and targets of 1,072–5,972 nodes
- **Meshes [52], [23]:** contains graphs from 3D object meshes, with motifs of 40–199 nodes and targets of 201–5,873 nodes; all motif–target pairs are unsatisfiable.

#### 2) Graph-database benchmark [57]

These datasets contain eight real-world graphs (see Table I), which are used as target graphs. Although the graphs are labeled, we ignore these labels in our experiments. The authors of [57] generated the pattern graphs as follows. For each target graph *G*′, pattern graphs are generated by randomly extracting connected subgraphs from *G*′ using a random walk strategy. For each target graph, nine pattern sets are constructed, each containing 200 connected pattern graphs with the same number of nodes, resulting in a total of 1,800 pattern instances per target graph. The sizes of the pattern graphs range from 4 to 20 nodes (i.e., 4, 8, 12, 16, and 20) for the Human and WordNet graphs, and from 4 to 32 nodes (i.e., 4, 8, 16, 24, and 32) for the other graphs. Except for pattern graphs of size 4, the query sets are divided into dense and sparse categories, with four dense and four sparse query sets for each target graph.

**TABLE I.** Attributes of the target graphs of the Graph-database benchmark. 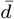 represents the average degree.

| Category | Graph | #nodes | #edges | $\bar{d}$ |
| --- | --- | --- | --- | --- |
| Biology | Yeast | 3,112 | 12,519 | 8.0 |
|  | Human | 4,674 | 86,282 | 36.9 |
|  | HPRD | 9,460 | 34,998 | 7.4 |
| Lexical | WordNet | 76,853 | 120,399 | 3.1 |
| Citation | US-Patents | 3,774,768 | 16,518,947 | 8.8 |
| Social | Youtube | 1,134,890 | 2,987,624 | 5.3 |
|  | DBLP | 317,080 | 1,049,866 | 6.6 |
| Web | Eu2005 | 862,664 | 16,138,468 | 37.4 |

#### 3) Scalability benchmark [47]

This benchmark contains three target graphs (see Table II) and is used for scalability experiments. The graphs hugetrace-00020 and road-usa are originally directed. Since Glasgow and PathLad+ do not support directed graphs, we converted these graphs into undirected graphs by treating each directed edge between two nodes as a single undirected edge. For each target graph *G*′, we generated pattern graphs by performing a random walk on *G*′. The generated pattern graphs are connected, and their sizes range from 9 to 499 nodes, covering 31 different sizes. For each size, we generated 50 pattern graph instances, resulting in a total of 1,550 instances per target graph.

**TABLE 2.**
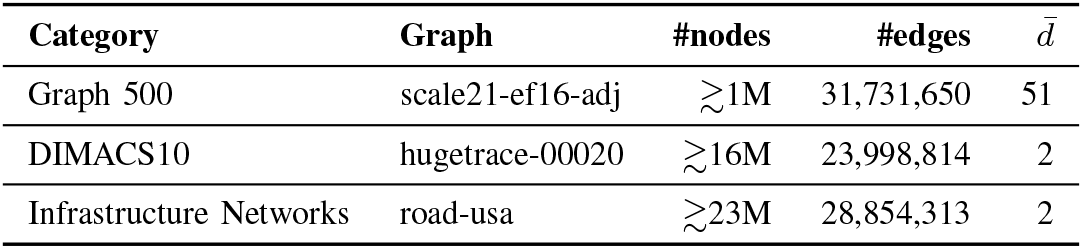
Attributes of the target graphs of the Scalability benchmark. 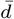 represents the average degree.

### B. Experiment setup

In our experiments, we measured both the matching time and memory usage required to solve the subgraph isomorphism problem. For instances from the Classic benchmark, we search for all solutions. For the Graph Database and Scalability benchmarks, the algorithms stop as soon as a solution is found, except for the hugetrace-00020 dataset, for which all solutions are searched. The default time limit is set to 5 hours for instances from the Classic benchmark (except for the MIVIA LDG dataset). If at least half of the solvers can solve a given instance within this limit, we attempt to solve the same instance with the remaining solvers using an extended timeout of 10 days. For the Graph-database benchmark, the timeout is set to 600 seconds, following previous works [57], [61]. For the Scalability benchmarks, the timeout is set to 1,200 seconds, i.e., doubled compared to the Graph-database benchmark, to account for the larger size of the graphs.

The experiments were conducted on a server equipped with an Intel(R) Xeon(R) Gold 6342 @ 2.80GHz processor with 54 cores, running in 64-bit mode, and featuring 1.7 MB of L1 cache, 216 MB of L2 cache, 864 MB of L3 cache, and 2 TB of RAM. The system was configured in NUMA mode with all 54 CPU cores assigned to a single NUMA node, ensuring optimized and balanced memory access for maximum performance. To accelerate the evaluation, we ran up to 50 instances in parallel for each algorithm, as all instances are independent and do not share data. This parallel execution strategy allows efficient utilization of the server resources without affecting the correctness or reproducibility of the results.

### C. What is the magnitude of the improvement achieved by FASTiso over VF3/VF3L?

To measure the improvement of FASTiso over VF3 and VF3L, we compare all three algorithms on the MIVIA LDG dataset, originally used by the authors of VF3 to demonstrate its superiority. We tested unlabeled graphs, graphs labeled according to a uniform distribution, and graphs labeled with a non-uniform distribution for all densities *d* (0.2, 0.3, 0.4). The average matching times for finding all solutions and for finding one solution for *d* = 0.3 is shown in Figure 1, all additional results are provided in the supplementary material. As shown by the results, Fastiso significantly outperforms VF3 across all datasets, and is outperformed by VF3L only on graphs labeled with a non-uniform distribution when enumerating all solutions.

**Fig. 1.**
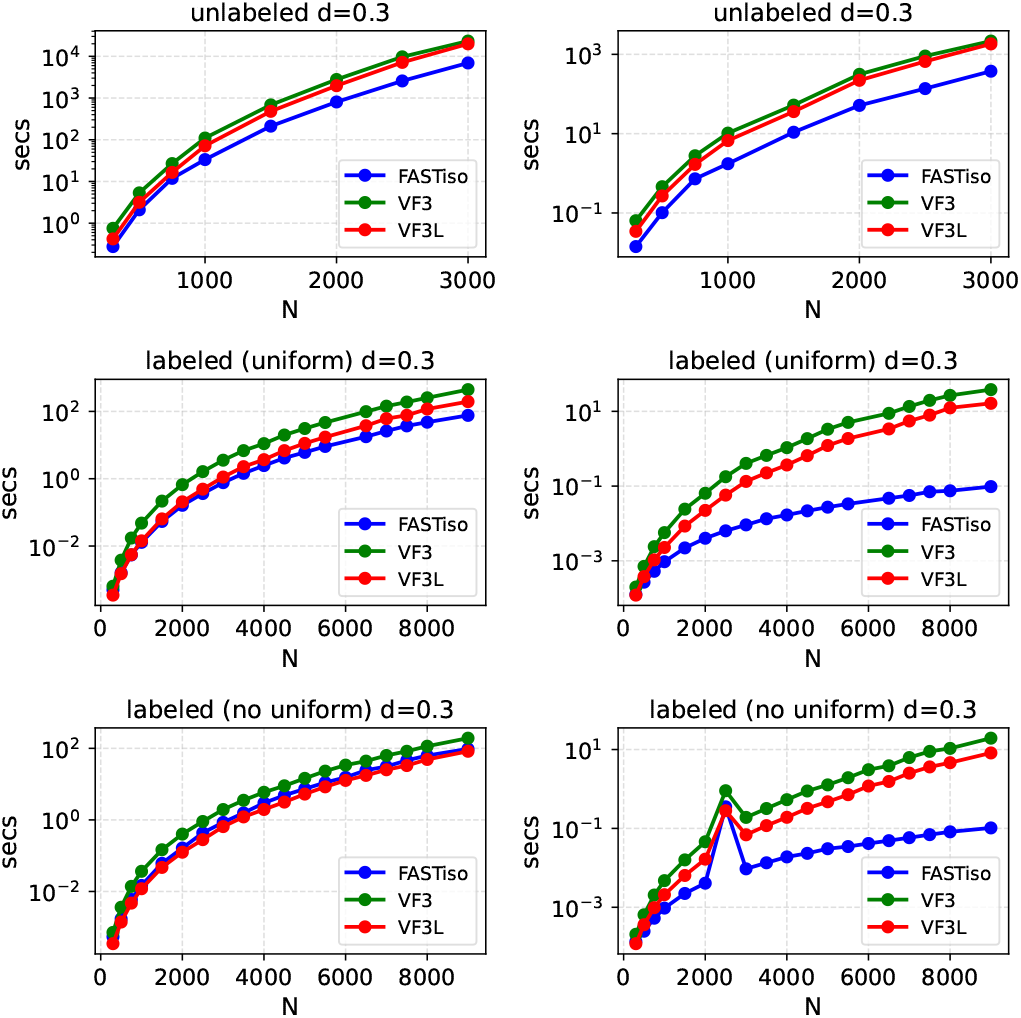
The average matching times for finding all solutions and for finding a single solution on graphs from the NIVIA LDG dataset with density *d* = 0.3. Here, *N* denotes the number of nodes in the target graph.

#### Finding all solutions

For unlabeled graphs, FASTiso’s advantage increases with both size (*N*) and density (*d*). It outperforms VF3L and VF3 with substantial time gains on large instances, notably for *d* = 0.3, *N* = 3000 (3–4 hours saved, speed-up of 2.87×–3.34×) and *d* = 0.4, *N* = 1500 (4–5 hours saved, up to 3.82× faster). For uniformly labeled graphs, FASTiso and VF3L exhibit similar performance for *d* = 0.2 and *N* ≤ 6500, both outperforming VF3. FASTiso becomes the fastest for *d* = 0.3, doubling VF3L’s speed for *d* = 0.4. In contrast, VF3L maintains a slight advantage (1.29×) on non-uniform label distributions.

#### Finding a single solution

FASTiso is consistently the fastest algorithm. For unlabeled graphs, it is at least twice as fast as its competitors, with a peak speed-up of 25.53× (*N* = 500, *d* = 0.4). For labeled graphs (uniform or non-uniform), it outperforms VF3 and VF3L by up to two orders of magnitude in certain sizes.

FASTiso also maintains a low memory consumption. With a 17% reduction in memory usage of FASTiso compared to VF3/VF3L. Detailed results on memory consumption are presented in the supplementary material.

### D. How does FASTiso compare with state-of-the-art solutions?

We compare FASTiso with VF3, VF3L, and the other state-of-the-art solvers on all Classic benchmarks, except for the MIVIA LDG dataset. The Figure 2 reports the detailed experimental results. Overall, FASTiso achieves the best global performance and is the fastest solver on the majority of the datasets (6 out of 9).

**Fig. 2.**
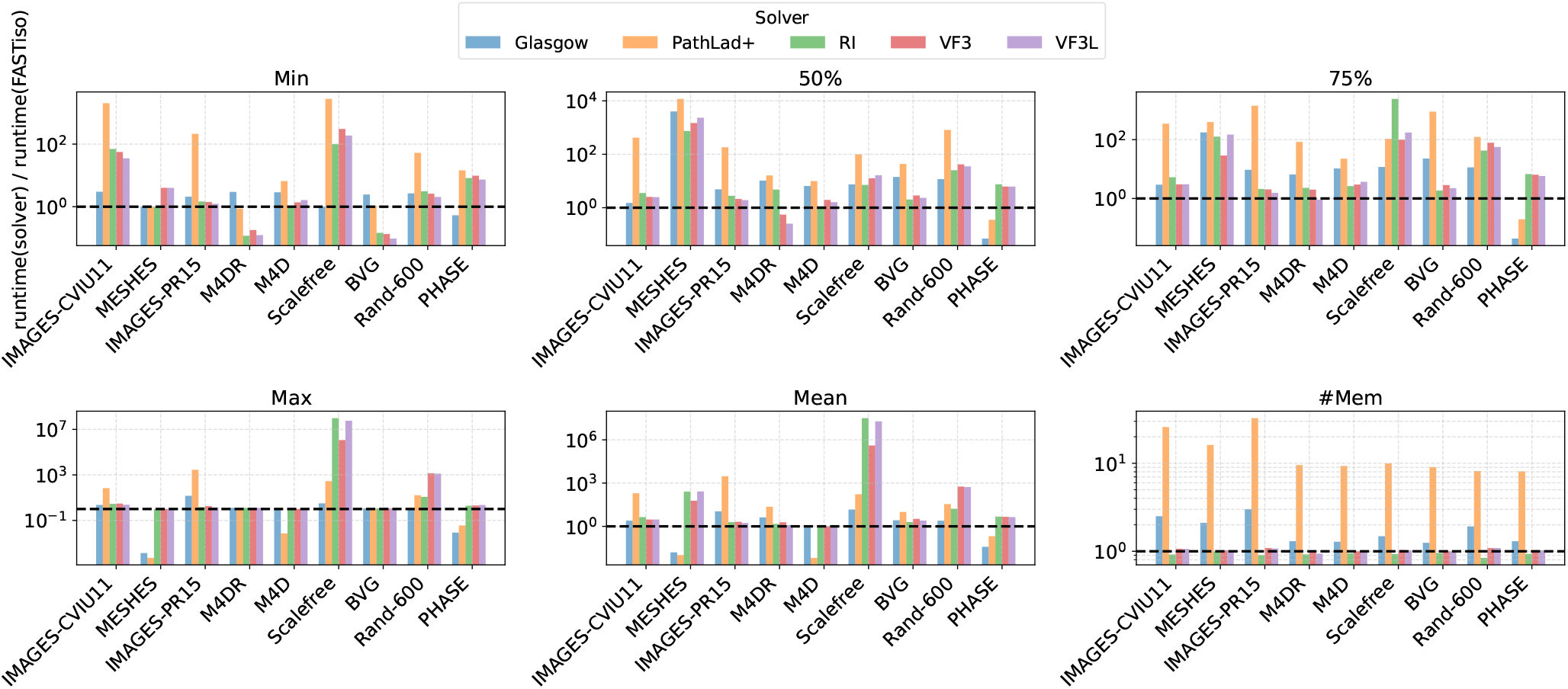
Detailed results on the Classic benchmark. We report the ratios of average runtime, maximum runtime, total runtime, and average memory usage of different solvers, normalized by the FASTiso solver (lower is better, FASTiso = 1). FASTiso systematically outperforms other tree search approaches and surpasses constraint programming-based methods on most datasets. However, FASTiso is mainly outperformed by constraint programming approaches on unsatisfiable instances, such as Phase and Meshes.

**Fig. 3.**
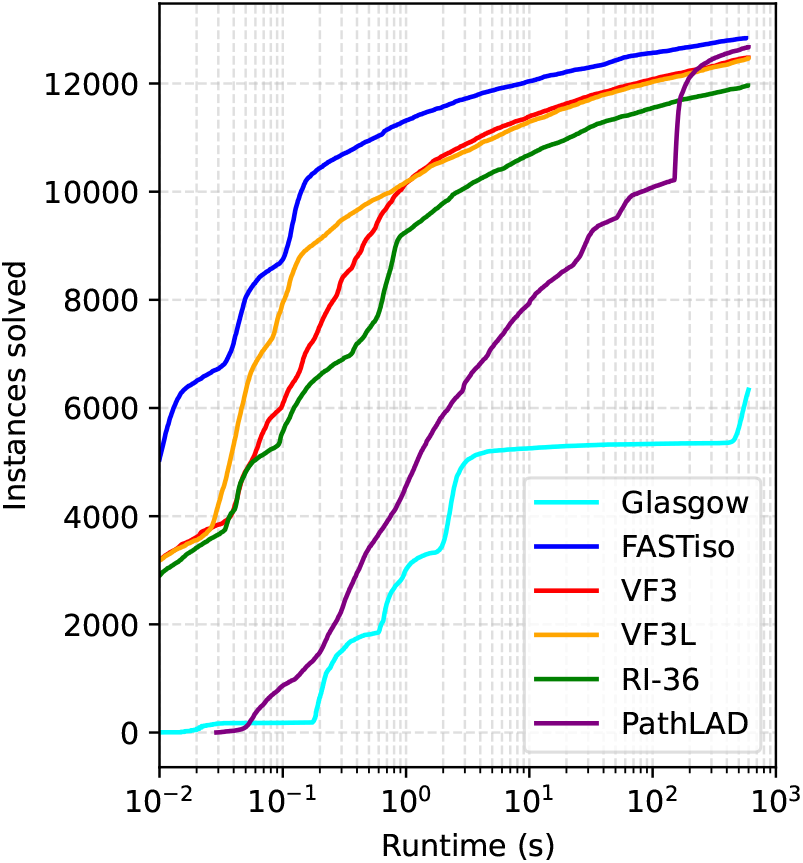
Detailed Results on Graph-database benchmark. We report the cumulative number of instances solved as a function of runtime.

Table 2 (see supplementary file) reports the results on the Scalefree dataset, which contains both sparse and dense graphs. This dataset is known to be particularly challenging for VF3 when compared to constraint programming-based approaches [14], [16]. As shown in the table, FASTiso achieves the best performance on this dataset, followed by Glasgow and PathLad+. Notably, PathLad+ is itself several orders of magnitude faster than VF3, VF3L, and RI. For most instances, all solvers exhibit similar execution times. However, FASTiso, Glasgow, and PathLad+ significantly widen the performance gap on a subset of instances corresponding to dense graphs.

These observations are further confirmed on the RAND dataset (see supplementary file, Table 3), which consists of random graphs, both sparse and dense, derived from the SI dataset. FASTiso is consistently the fastest solver across all graph sizes. It is approximately twice as fast as Glasgow, and outperforms RI, VF3, VF3L, and PathLad+ by one to two orders of magnitude for graph sizes 200 and 600.

Overall, FASTiso performs particularly well on small dense graphs. An exception is observed on the Phase dataset (see supplementary file, Table 10), which consists of very dense graphs of small size. This dataset contains only unsatisfiable instances, where advanced filtering and constraint propagation techniques give Glasgow and PathLad+ a distinct advantage. On this dataset, Glasgow is the fastest solver, followed by PathLad+ and FASTiso. Glasgow outperforms FASTiso by approximately one order of magnitude, while FASTiso remains about four times faster than VF3, VF3L, and RI.

The six remaining datasets of the classic benchmark contain only small and sparse graphs. Even on these datasets, FASTiso exhibits the best overall performance, ranking as the top solver on four out of six datasets. It consistently outperforms VF3, VF3L, and RI, but is outperformed by Glasgow and PathLad+ on the Meshes dataset, which, like the Phase dataset, consists exclusively of unsatisfiable instances. However, the superiority of Glasgow and PathLad+ over FASTiso is observed on only a single instance that FASTiso fails to solve within the allotted time. Finally, FASTiso is also outperformed by PathLad+ on the M4D dataset.

Table III and Figure 2 report the results of our experiments on the Graph-database benchmark datasets. Glasgow fails to solve any instance on several datasets due to the high computational complexity of its filtering techniques, and consumes more than 500 GB of memory on each instance. These datasets are reported as NA in Table III. FASTiso solves the largest number of instances on five out of the eight datasets. Overall, FASTiso solves 12,837 instances, followed by PathLAD+ with 12,670 instances, VF3 with 12,477 instances, VF3L with 12,463 instances, and Glasgow with 6,336 instances. The most challenging dataset for all solvers is Eu2005, which contains the largest number of edges. However, it can be observed that constraint programming–based approaches are more effective than tree-search–based approaches on the Human dataset.

**TABLE 3.** Comparison on the Graph-database benchmark. Each dataset contains 1,800 instances. For each solver, we report the number of instances solved.

| solver | Yeast | Human | HPRD | WordNet | DBLP | Eu2005 | Youtube | US-Patents |
| --- | --- | --- | --- | --- | --- | --- | --- | --- |
| FASTiso | 1744 | 1193 | 1794 | **1800** | **1730** | **1161** | **1732** | **1683** |
| VF3 | 1720 | 1084 | 1786 | 1795 | 1712 | 1096 | 1637 | 1647 |
| VF3L | 1724 | 1043 | 1782 | 1794 | 1696 | 1131 | 1636 | 1657 |
| RI | 1727 | 950 | 1713 | 1796 | 1481 | 1011 | 1637 | 1646 |
| PathLad | **1798** | 1757 | 1798 | 1771 | 1630 | 1047 | 1688 | 1181 |
| Glasgow | 1797 | **1763** | **1800** | 976 | NA | NA | NA | NA |

### E. Scalability Test

Figures 4 and 5 report the scalability results. The Glasgow solver is not included in these figures, as it requires more than 500 GB of memory, and the time complexity of its filtering and constraint propagation techniques prevents it from solving the large instances of this benchmark.

**Fig. 4.**
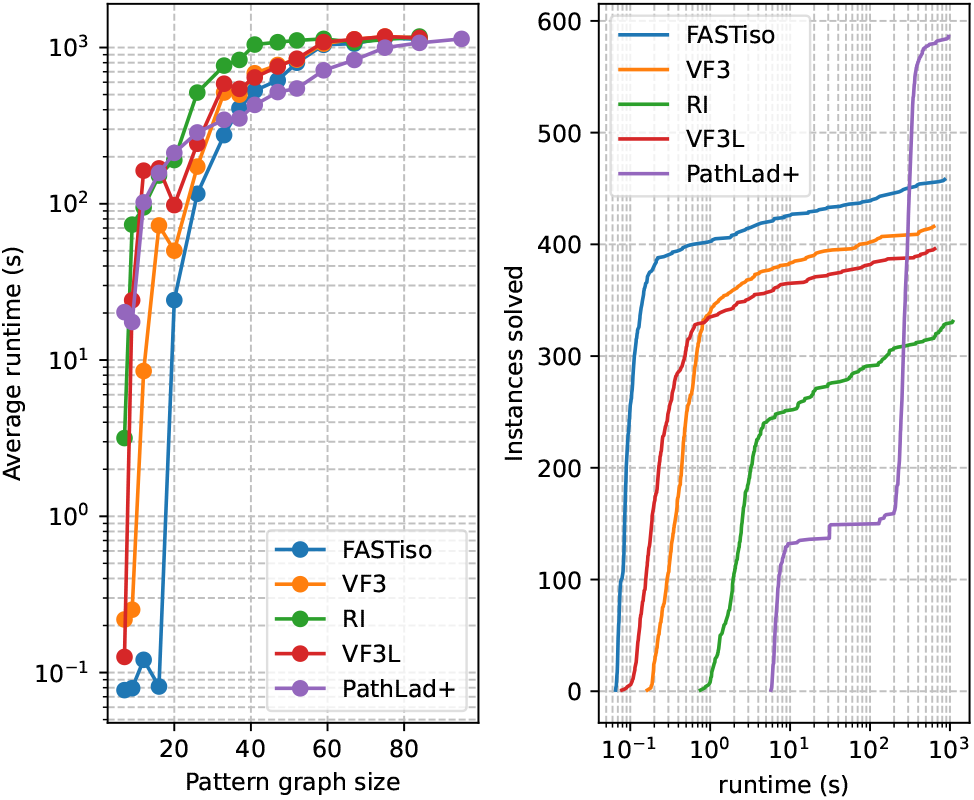
Detailed Results on scale21-ef16-ad dataset (Scalability benchmark). On the left, we report the mean runtime with respect to the pattern graph size in terms of number of nodes, and on the right, the cumulative number of instances solved as a function of runtime. The peak memory usage of FASTiso is 5.15 GB, whereas PathLAD+ reaches 4.17GB.

**Fig. 5.**
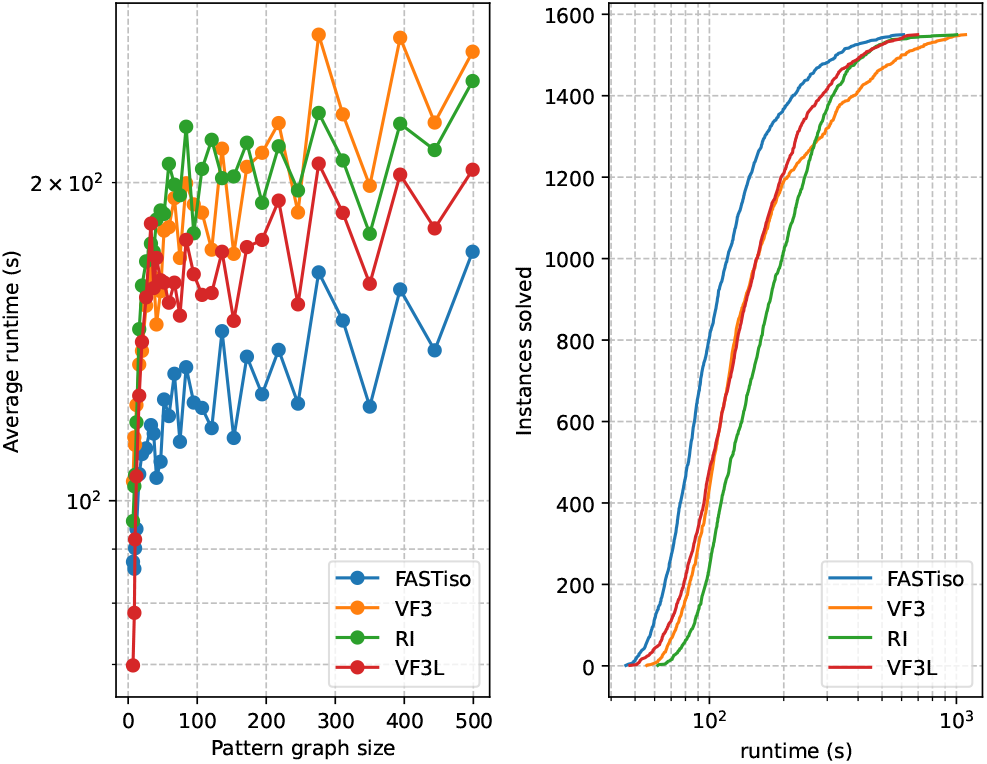
Detailed Results on hugetrace-00020 dataset (Scalability benchmark). On the left, we report the mean runtime with respect to the pattern graph size in terms of number of nodes, and on the right, the cumulative number of instances solved as a function of runtime. PathLAD+ does not solve any instance in this dataset. The peak memory usage of FASTiso is 7.74 GB, whereas PathLAD+ reaches 36.19 GB.

As observed for the Eu2005 dataset in the Graph-database benchmark, the dataset on which the solvers resolve the fewest instances is scale21-ef16-adj, which also has the largest number of edges. This indicates a clear sensitivity of the algorithms to the number of edges. On the scale21-ef16-adj dataset, PathLad+ solves the largest number of instances (586), followed by FASTiso (458), VF3 (416), VF3L (396), and RI (331). Although FASTiso is faster than PathLad+ for pattern graphs of size approximately smaller than 40, the more sophisticated filtering and constraint propagation techniques employed by PathLad+ allow it to outperform FASTiso for pattern graphs of size greater than or equal to 40.

However, this advantage does not hold on the hugetrace-00020 dataset, whose target graph contains more than 16 million nodes. On this dataset, PathLad+ does not solve any instance within the time limit, whereas FASTiso and the other solvers successfully solve all instances. This suggests that PathLad+ is more sensitive to the number of nodes than FASTiso. To further confirm this observation, we compare FASTiso and PathLad+ on the road-usa dataset, whose target graph contains more than 23 million nodes. The results, reported in the supplementary material (see Figure D), are consistent with those obtained on hugetrace-00020: PathLad+ solves 275 instances, while FASTiso solves 1358 instances.

Although FASTiso is not the best-performing algorithm on the scale21-ef16-adj dataset, its new pruning rules, which are more effective than those of other tree search methods and less costly than those used by PathLad+ and Glasgow, allow it to outperform the other tree search methods on the scale21-ef16-adj and hugetrace-00020 datasets.

## V. Conclusion

In this article, we propose FASTiso, an exact subgraph isomorphism algorithm that emphasizes a strong consistency between the variable ordering strategy and the pruning rules used during the mapping process. Extensive experiments conducted on a wide range of reference datasets demonstrate the effectiveness and robustness of FASTiso. The results show that FASTiso adapts well to diverse graph structures, performing efficiently on both small and large graphs. Overall, it achieves the best global performance among the evaluated algorithms and clearly outperforms its reference solver VF3, as well as the other tree-search-based methods such as RI and VF3L, on the vast majority of datasets. The only exception is observed on the MIVIA LDG dataset, where VF3L slightly outperforms FASTiso on graphs labeled with a non-uniform distribution when enumerating all solutions.

Compared to constraint programming-based approaches, FASTiso achieves competitive or superior performance on most datasets, while being outperformed on a few instances where advanced filtering and constraint propagation techniques provide a significant advantage. However, due to the higher computational overhead of these techniques, constraint-based solvers exhibit a stronger sensitivity to the number of nodes. In contrast, FASTiso demonstrates better scalability behavior, making it more suitable for large-scale graphs.

Looking ahead, we plan to further improve FASTiso by exploring additional pruning rules and incorporating symmetry handling techniques to reduce redundant exploration. We also intend to develop a parallel and distributed version of the algorithm to handle very large graphs containing billions of nodes.

## Supporting information

Supp. mat.

