## Supplementary material for "FASTiso: Fast Algorithm on Search state Tree for subgraph ISOmorphism in graphs of any size and density": Supp. mat.

### I. Supplementary material

#### A. Appendix

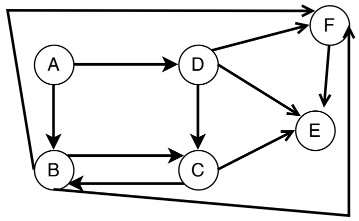


| Node | A | B | C | D | E | F | $\mu_G$ |
|---|---|---|---|---|---|---|---|
| Deg | 2 | 5 | 4 | 4 | 3 | 4 | |
| Step 0 | | | | | | | |
| | | | | | | | $\emptyset$ |
| Step 1 | | | | | | | |
| $\lvert N_1(u_i)\rvert$ | $\lvert\{B\}\rvert$ | - | $\lvert\{B\}\rvert$ | 0 | 0 | $\lvert\{B\}\rvert$ | $\{B\}$ |
| $N_{\text{edges}}(u_i, N_1(u_i))$ | 1 | - | 2 | 0 | 0 | 2 | |
| $\sum_{u_a \in N_2(u_i)} \lvert N_1(u_a)\rvert$ | 0 | - | 0 | 3 | 2 | 0 | |
| $\lvert N_3(u_i)\rvert$ | $\lvert\{D\}\rvert$ | - | $\lvert\{D,E\}\rvert$ | $\lvert\{E\}\rvert$ | $\lvert\{D\}\rvert$ | $\lvert\{D,E\}\rvert$ | |
| Step 2 | | | | | | | |
| $\lvert N_1(u_i)\rvert$ | $\lvert\{B\}\rvert$ | - | - | $\lvert\{C\}\rvert$ | $\lvert\{C\}\rvert$ | $\lvert\{B\}\rvert$ | $\{B,C\}$ |
| $N_{\text{edges}}(u_i, N_1(u_i))$ | 1 | - | - | 1 | 1 | 2 | |
| $\sum_{u_a \in N_2(u_i)} \lvert N_1(u_a)\rvert$ | 1 | - | - | 3 | 2 | 2 | |
| $\lvert N_3(u_i)\rvert$ | 0 | - | - | 0 | 0 | 0 | |

Fig. A. Computation of the variable ordering for the graph on the left using our strategy. For simplicity, we assume that all nodes have the same initial domain; therefore, this information is not discriminative for computing the variable ordering. Only the first two steps are shown due to space constraints.

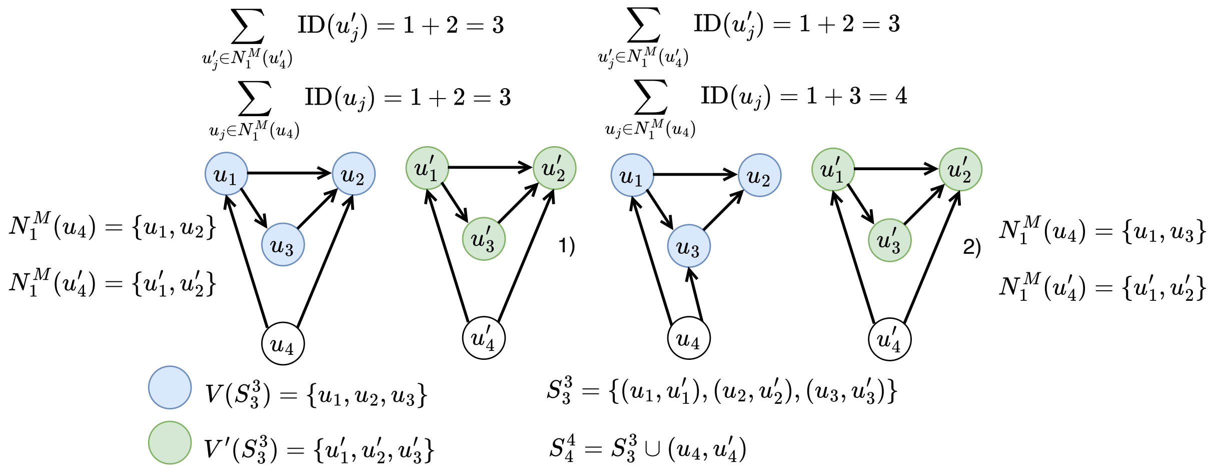


Fig. B. Verification of isomorphism conditions when adding the pair $(u_4, u'_4)$ to state $S_3^3$. In Figure A), this addition results in a new consistent state $S_4^4$. In contrast, in Figure B), the addition does not lead to a new consistent state. Although the first three conditions of $F_c$ are satisfied, the condition given by Equation (3) is not satisfied.

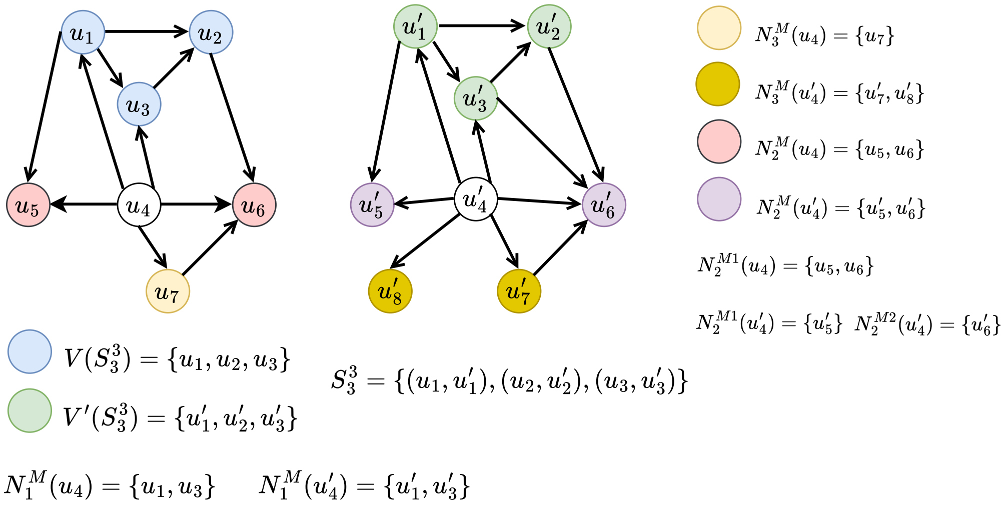


Fig. C. Verification of a dead state when adding the pair $(u_4, u'_4)$ to state $S_3^3$. This addition leads to a new consistent state; however, this state can never evolve into a goal state. Indeed, although the first two conditions of $F_d$ are satisfied, the condition given by Equation (4) is violated, since $N_2^{M_1}(u_4) = 2 > N_2^{M_1}(u'_4) = 1$.

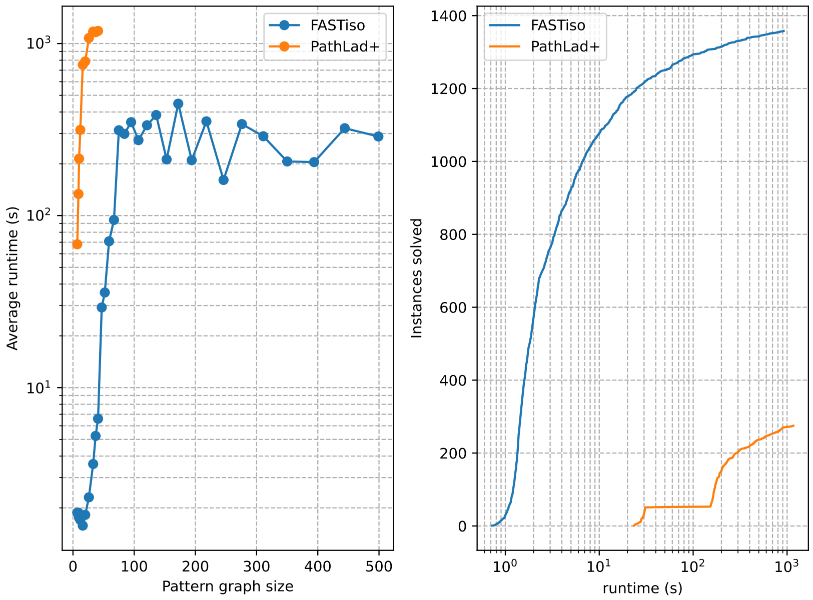


Fig. D. Detailed Results on road-usa dataset (Scalability benchmark). On the left, we report the mean runtime with respect to the pattern graph size in terms of number of nodes, and on the right, the cumulative number of instances solved as a function of runtime.

| Class | #inst | P. Nodes Min | P. Nodes Max | P. Edges Min | P. Edges Max | P. Dens. Min | P. Dens. Max | T. Nodes Min | T. Nodes Max | T. Edges Min | T. Edges Max | T. Dens. Min | T. Dens. Max |
|---|---|---|---|---|---|---|---|---|---|---|---|---|---|
| **MIVIA LDG** | | 60 | 2000 | 708 | 1599200 | 0.2 | 0.4 | 300 | 10000 | 17940 | 39996000 | 0.2 | 0.4 |
| **SF** | 100 | 180 | 900 | 956 | 11956 | 0.01 | 0.17 | 200 | 1000 | 1184 | 14296 | 0.01 | 0.16 |
| **RAND** | 270 | 40 | 360 | 82 | 24820 | 0.02 | 0.21 | 200 | 600 | 872 | 68420 | 0.02 | 0.20 |
| **Phase** | 50 | 30 | 30 | 256 | 424 | 0.29 | 0.49 | 150 | 150 | 8264 | 11842 | 0.37 | 0.53 |
| **BVG** | 540 | 40 | 480 | 86 | 4274 | 0.01 | 0.2 | 200 | 800 | 598 | 7200 | 0.00 | 0.05 |
| **M4D** | 90 | 51 | 777 | 192 | 4116 | 0.00 | 0.08 | 256 | 1296 | 1344 | 7200 | 0.00 | 0.02 |
| **M4DR** | 270 | 51 | 777 | 152 | 4150 | 0.00 | 0.08 | 256 | 1296 | 1444 | 8757 | 0.00 | 0.03 |
| **Images-PR15** | 24*1 | 4 | 170 | 8 | 483 | 0.02 | 0.67 | 4838 | 4838 | 14134 | 14134 | 0.00 | 0.00 |
| **Images-CVIU11** | 43*146 | 15 | 151 | 40 | 430 | 0.02 | 0.19 | 1072 | 5972 | 3080 | 17782 | 0.00 | 0.00 |
| **Meshes** | 6*503 | 40 | 199 | 228 | 1078 | 0.02 | 0.15 | 201 | 5873 | 504 | 30585 | 0.00 | 0.02 |

Fig. E. Attributes of the graphs of the Classic benchmarks datasets.

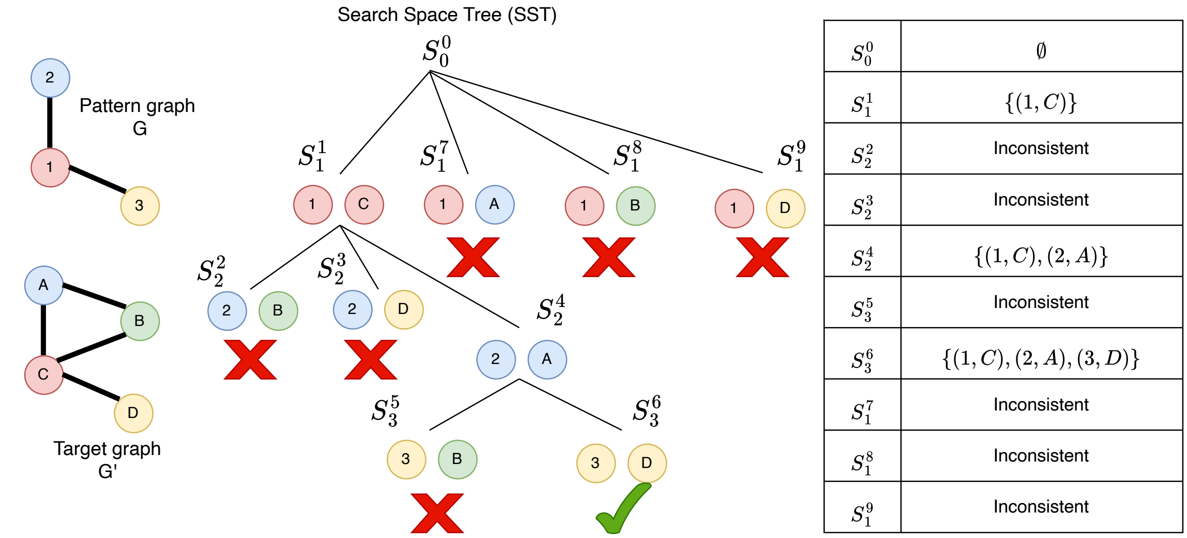


Fig. F. Search space tree. Letters and numbers on the nodes represent their identifiers. Nodes of the pattern graph can only be matched with nodes of the target graph having the same color. State $S_3^6$ is a goal state since it satisfies all subgraph isomorphism conditions and is complete ($|V(G)| = 3$). Inconsistent states are marked with red crosses; their inconsistency results from violating the node label preservation constraint (Condition 5).

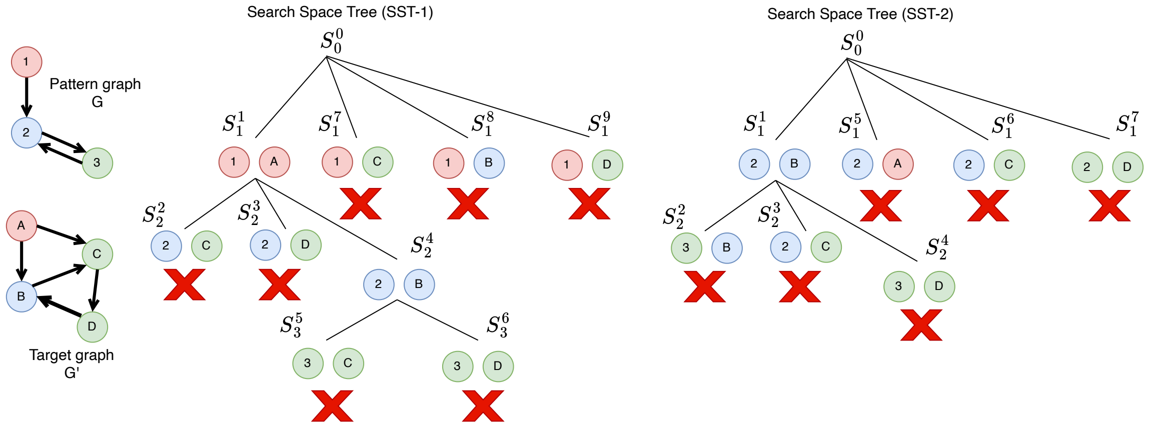


Fig. G. Impact of variable ordering on the search space tree. Letters and numbers on the nodes represent their identifiers. Nodes of the pattern graph can only be matched with nodes of the target graph having the same color. SST-1 is the search space tree induced by the static variable ordering 1-2-3, which selects first the nodes of $G$ with the rarest labels in $G'$. SST-2 is the search space tree induced by the static variable ordering 2-3-1, which prioritizes the nodes of $G$ with the highest degree. In case of ties (nodes having the same degree), the node whose label is the rarest in $G'$ is selected. As illustrated, the ordering 2-3-1 is more effective, as it generates fewer states and explores a smaller search space than the ordering 1-2-3.

TABLE 1
Notation used in the paper. Let $G = (V, E)$ be a pattern graph and $G' = (V', E')$ be a target graph.

| Symbol | Meaning |
|---|---|
| $N^+(u)$ | Out-neighborhood of node $u$: $N^+(u) = \{v \in V \mid (u, v) \in E\}$ |
| $N^-(u)$ | In-neighborhood of node $u$: $N^-(u) = \{v \in V \mid (v, u) \in E\}$ |
| $N(u)$ | Total neighborhood of $u$: $N(u) = N^+(u) \cup N^-(u)$ |
| $d^+(u)$ | Out-degree of $u$: number of outgoing edges (counting multiplicities) |
| $d^-(u)$ | In-degree of $u$: number of incoming edges (counting multiplicities) |
| $d(u)$ | Total degree of $u$: $d(u) = d^+(u) + d^-(u)$ |
| $N_{\text{edges}}(u_i, S)$ | Total number of edges (incoming and outgoing) between node $u_i$ and nodes in $S$, counting multiplicities |
| $N_{\text{in-edges}}(u_i, S)$ | Total number of incoming edges between node $u_i$ and nodes in $S$, counting multiplicities |
| $N_{\text{out-edges}}(u_i, S)$ | Total number of outgoing edges between node $u_i$ and nodes in $S$, counting multiplicities |
| $S_k^c$ | Partial state (mapping) at step $k$ in the SST: set of pairs $(u_i, M(u_i)), i < k$ |
| $V(S_k^c)$ | Nodes of $G$ included in $S_k^c$: $\{u_i \mid (u_i, M(u_i)) \in S_k^c\}$ |
| $V'(S_k^c)$ | Nodes of $G'$ included in $S_k^c$: $\{M(u_i) \mid (u_i, M(u_i)) \in S_k^c\}$ |
| $\mu_G$ | Variable ordering of the nodes in $G$ |
| $\mu_G(k)$ | Partial variable ordering after $k$ nodes have been selected |
| $N_1(u_i)$ | Neighbors of $u_i$ already included in $\mu_G(k)$: $\{u_j \in N(u_i) \mid u_j \in \mu_G(k)\}$ |
| $N_2(u_i)$ | Neighbors of $u_i$ not in $\mu_G(k)$ but adjacent to at least one node in $\mu_G(k)$: $\{u_j \in N(u_i) \mid u_j \in N(\mu_G(k))\}$ |
| $N_3(u_i)$ | Neighbors of $u_i$ not in $N_1(u_i)$ nor $N_2(u_i)$ |
| $N_1^M(u)$ | Set of neighbors of $u$ that are already mapped |
| $N_2^M(u)$ | Set of neighbors of $u$ not yet mapped but having neighbors already mapped |
| $N_3^M(u)$ | Set of neighbors of $u$ that belong neither to $N_1^M(u)$ nor $N_2^M(u)$ |
| $N_2^{Mi}(u)$ | Subset of $N_2^M(u)$ containing unmapped neighbors of $u$ with exactly $i$ mapped neighbors $(i > 0)$ |

##### B. Variable ordering

This section discusses a scalability issue related to our variable ordering strategy and proposes a solution.

One way to efficiently implement the variable ordering strategy is to update node information incrementally. This approach leads in the worst case to $O\big(|V|^2 + \sum_{u_i \in V} |N_1^M(u_i)| \cdot |N(u_i)|\big)$, where $N_1^M(u_i)$ denotes the number of mapped neighbors of $u_i$ at the time it is selected.

For sparse graphs, the number of neighbors and mapped neighbors remains limited, and the resulting complexity tends toward $O(|V|^2 + |E|)$, which is of the same order as the variable ordering strategies used in RI and VF3, and therefore does not raise scalability concerns.

In contrast, for very dense graphs, both the number of neighbors and the number of mapped neighbors may become proportional to $|V|$. In this case, the term $\sum_{u_i \in V} |N_1(u_i)| \cdot |N(u_i)|$ may grow up to $O(|V|^3)$, leading to a prohibitive computational cost and limiting scalability. To address this issue, we introduce a parameter $p$ that bounds the number of mapped neighbors considered during the evaluation. Specifically, when assessing the influence of a node, at most $p$ mapped neighbors are taken into account. This bounding modifies the complexity to $O\big(|V|^2 + \sum_{u_i \in V} p \cdot |N(u_i)|\big)$. When $p$ is constant, the complexity simplifies to $O(|V|^2 + |E|)$, thereby eliminating the cubic behavior observed on dense graphs and significantly improving scalability.

Let $\mu_G$ denote the variable ordering obtained without bounding, and $\mu_G^p$ the ordering obtained with parameter $p$. The first $p$ nodes in $\mu_G$ and $\mu_G^p$ are identical, while the ordering of the remaining nodes may differ. Although this approximation may slightly affect search efficiency, it provides a favorable trade-off between computational cost and ordering quality. Moreover, if the maximum degree of $G$ is smaller than $p$, then $\mu_G = \mu_G^p$.

In our implementation of FASTiso, we set $p = 50$.

##### C. Implementation details of the component $F_c$

The use of filtering tests to avoid verifying condition 4 of $F_c(S_k^c, u_k, u_k')$ as much as possible will only be effective if these tests have a complexity lower than that of verifying condition 4. We will present the implementation details that allow these filtering tests to be checked in constant time $\Theta(1)$.

To manage the sets $V(S_k^c)$ and $V'(S_k^c)$, two lists are used, structured as follows: $M[\mathrm{ID}(u_k)] = u_k'$ if $u_k$ is mapped to $u_k'$, and null otherwise (see Algorithm 1, Line 1-2). These structures make it possible to find the correspondent of a node and verify the membership of this node in $V(S_k^c)$ and $V'(S_k^c)$ in $\Theta(1)$.

Filtering tests are also performed in $\Theta(1)$ using lists that store, for each node, the relevant information required for filtering: the number of mapped neighbors (num_mapped_neighbors), the number of incoming edges with mapped neighbors (num_in_mapp), the number of outgoing edges with mapped neighbors (num_out_mapp), and the sum of the identifiers of the mapped neighbors (sum_ids).

Since FASTiso uses a static variable ordering, at each level $k$ of the SST corresponds the same node of the pattern graph $u_k = \mu(k)$. These lists are therefore precomputed before the matching process for the nodes of the pattern graph. For the nodes of the target graph, these lists are updated when a pair $(u_k, u_k')$ is added or removed, in $\Theta(|N(u_k')|)$ (see Algorithm 1).

Algorithm 1: addPair: Updates the current state $S_k^c$ by adding the pair $(u_k, u_k')$. The mapping structures and filtering information for neighboring nodes are updated accordingly. The function removePair($S_k^c$,$u_k$,$u_k'$) performs the inverse operation of addPair. It decrements the corresponding values in $S_k^c$ and sets $M[\mathrm{ID}(u_k)] = M'[\mathrm{ID}(u_k')] = \mathrm{null}$

Data: $S_k^c$,$u_k$,$u_k'$

1 $M[ID(u_k)] \leftarrow u_k'$;
2 $M'[ID(u_k')] \leftarrow u_k$;
3 for $u_j'$ in $N(u_k')$ do
4     num_mapped_neighbors[$ID(u_j')$]+=1;
5     num_in_mapp[$ID(u_j')$]+=$N_{\text{in-edges}}(u_k', \{u_j'\})$;
6     num_out_mapp[$ID(u_j')$]+=$N_{\text{out-edges}}(u_k', \{u_j'\})$;
7     sum_ids[$ID(u_j')$]+=$ID(u_k)$
8 return $S_k^c \cup \{(u_k, u_k')\}$;

##### D. Implementation details of the component $F_d$

To manage the classified neighbor sets for a node $u$, a list $L$ of size $t = \max_{v \in N_2^M(u)} |N_1^M(v)|$ is used, where the value at an index $i > 0$ corresponds to $|N_2^{Mi}(u)|$, and the value at index $i = 0$ corresponds to $|N_3^M(u)|$. This configuration allows for an easy calculation of $|N_2^M(u)| = |N(u)| - |N_1^M(u)| - |N_3^M(u)|$ and the verification of the last condition of $F_d(S_k^c, u_k, u'_k)$ in $\Theta(t)$. For nodes $u_k$ in the pattern graph, $L$ is precomputed before the matching process. However, for a node $u'_k$ in the target graph, $L$ is computed during the addition of the pair $(u_k, u'_k)$ to $S_k^c$ in the function $F_d(S_k^c, u_k, u'_k)$. This computation is performed in $\Theta(N(u'_k))$ by iterating over the neighbors $u'_j$ of $u'_k$ and incrementing $L[|N_1^M(u'_j)|]$ by 1.

##### E. Non-induced subgraph and graph isomorphism variants of isFeasible($S_k^c$, $u_k$, $u'_k$)

We have also proposed versions for graph isomorphism and monomorphism (non-induced subgraph isomorphism) for isFeasible($S_k^c$, $u_k$, $u'_k$)

For every pair $(u_k, u'_k)$ to be added to $S_k^c$:

Graph Isomorphism: For graph isomorphism, $F_c(S_k^c, u_k, u'_k)$ verifies the same conditions as for subgraph isomorphism. Additionally, $F_d(S_k^c, u_k, u'_k)$ must satisfy:

$$\begin{aligned} |N_2^M(u_k)| &= |N_2^M(u'_k)|, \\ |N_3^M(u_k)| &= |N_3^M(u'_k)|, \\ |N_2^{Mi}(u_k)| &= |N_2^{Mi}(u'_k)| \quad \forall\, 1 < i < \text{neighbs\_dm}[k]. \end{aligned} \tag{I}$$

Monomorphism: For monomorphism, $F_c(S_k^c, u_k, u'_k)$ must satisfy:

$$\begin{aligned} u'_k &\notin V'(S_k^c), \\ N_1^M(u_k) &\leq N_1^M(u'_k), \\ N_{\text{in}}(u_k) &\leq N_{\text{in}}(u'_k), \\ N_{\text{out}}(u_k) &\leq N_{\text{out}}(u'_k), \\ \forall \{u_k, u_a\} \in E,\ u_a \in V(S_k^c), &\ \exists \{u'_k, M(u_a)\} \in E' \\ &\text{s.t. } \beta_G(\{u_k, u_a\}) = \beta_{G'}(\{u'_k, M(u_a)\}). \end{aligned} \tag{II}$$

Additionally, $F_d(S_k^c, u_k, u'_k)$ must verify:

$$\begin{aligned} |N_2^M(u_k)| &\leq |N_2^M(u'_k)|, \\ \max_{u_j \in N_2^M(u_k)} |N_1^M(u_j)| &\leq \max_{u'_j \in N_2^M(u'_k)} |N_1^M(u'_j)|, \\ \sum_{u_j \in N_2^M(u_k)} |N_1^M(u_j)| &\leq \sum_{u'_j \in N_2^M(u'_k)} |N_1^M(u'_j)|. \end{aligned} \tag{III}$$

##### F. Proofs of correctness and exhaustiveness

We will first show that any solution returned by FASTiso is an isomorphism, then that the lookahead performed by isFeasible to detect dead states should never reject a pair $(u_k, u'_k)$ if its addition will result in a non-dead state, and finally that any valid isomorphism is returned by FASTiso.

Lemma 1: If FASTiso returns a solution, then it is valid.

Proof: By construction every time a matching $(u_k, u'_k)$ is added to a partial solution, isFeasible checks whether the isomorphism properties are preserved (see Section III-D. Therefore any solution returned by FASTiso is necessarily valid.

Lemma 2: isFeasible never reject a pair $(u_k, u'_k)$ if its addition will result in a non-dead state.

Proof: At a level $k$ of the search tree, a node $u_k$ can only be matched to a node $u'_k$ if they have the same number of already mapped neighbors (this condition is necessary but not sufficient): $|N_1^M(u_k)| = |N_1^M(u'_k)|$.

We show that if $|N_1^M(u_k)| \neq |N_1^M(u'_k)|$ at a level $l$ of the search tree, then $u_k$ and $u'_k$ can never be matched at a level $k > l$.

Suppose that $u_k$ and $u'_k$ are matched at level $k$, but at some level $l < k$ we had $|N_1^M(u_k)| \neq |N_1^M(u'_k)|$. For $|N_1^M(u_k)| = |N_1^M(u'_k)|$ to become true at level $k$, each new matching $(u, u')$ added between levels $l$ and $k-1$ must modify $N_1^M(u_k)$ and $N_1^M(u'_k)$ differently. This can only happen if a neighbor $u$ of $u_k$ is matched to a node $v$ that is not a neighbor of $u'_k$, and vice versa, which contradicts the subgraph isomorphism constraints.

It follows that $u_k$ and $u'_k$ cannot be mapped together, contradicting our assumption. Thus, if $|N_1^M(u_k)| \neq |N_1^M(u'_k)|$ at level $l$, then they can never be matched at a level $k > l$.

The function isFeasible exploits this property to anticipate dead states. Specifically, for each pair $(u_k, u'_k)$, it classifies the unmapped neighbors of $u_k$ and $u'_k$ based on the number of already mapped neighbors. Then, it checks whether a classified set of neighbors of $u_k$ is larger than its equivalent for $u'_k$. If so, it is known that some neighbors of $u_k$ will never find a correspondent at a level $l > k$ of the search tree.

Thus, the function isFeasible never reject a pair $(u_k, u'_k)$ if its addition will result in a non-dead state.

Lemma 3: All valid solutions are returned by FASTiso

Proof: FASTiso uses exhaustive backtracking, ensuring that it explores all possible correspondences between $G$ and $G'$, subject to the conditions imposed by the function isFeasible.

Suppose that there exists a valid isomorphism $M$ that is not found by FASTiso. This implies that at some level $k$ of the search tree, the function isFeasible rejects a matching $(u_k, u'_k)$ belonging to $M$. However, we know that the function isFeasible never reject a pair $(u_k, u'_k)$ if its addition will result in a non-dead state (Lemma 2). This leads to a contradiction, proving that FASTiso necessarily returns all valid solutions.

#### G. How do the new variable ordering and pruning rules introduced by FASTiso influence their efficiency?

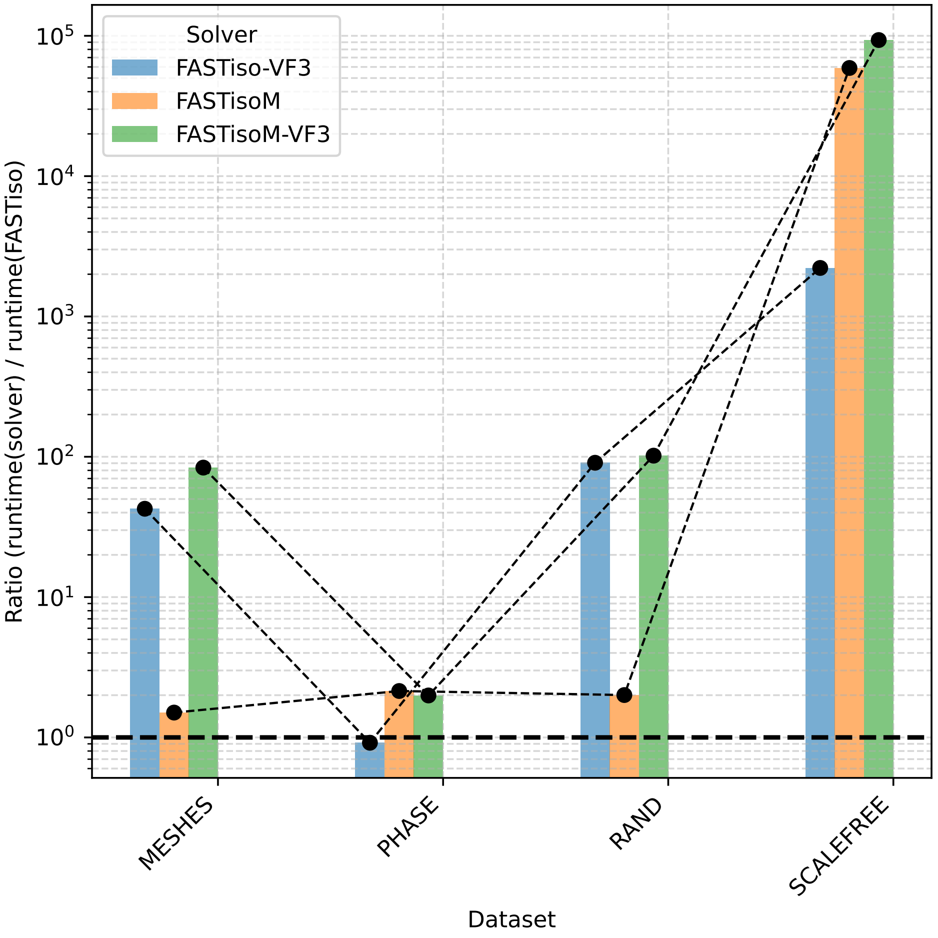


Fig. H. Ablation study of FASTiso on four representative datasets. We report the runtime ratios of different FASTiso variants normalized by the full FASTiso solver (lower is better, FASTiso = 1). FASTiso-VF3 removes the proposed variable ordering and uses the VF3 ordering instead, FASTisoM removes the two novel pruning rules; FASTisoM-VF3 removes both contributions and relies on the VF3 ordering.

To assess the effectiveness of the proposed variable ordering strategy and the two novel pruning rules, we conduct an ablation study by comparing several variants of FASTiso against the full solver. The evaluation is performed on four representative datasets: SCALEFREE, RAND, Meshes, and Phase. Due to time constraints, we deliberately selected two datasets where FASTiso achieves the best performance (SCALEFREE and RAND) and two datasets where it is outperformed by at least one competing solver (Meshes and Phase). We consider the following variants: 1–) FASTiso-VF3, which corresponds to FASTiso without the proposed variable ordering and instead relies on the variable ordering strategy of VF3; 2–) FASTisoM, which removes the two proposed pruning rules while keeping the new variable ordering; and 3–) FASTisoM-VF3, which disables both the proposed variable ordering and the pruning rules, and uses the VF3 variable ordering. For each variant, we report the runtime ratio with respect to the full FASTiso solver, where lower values indicate better performance and the full FASTiso is normalized to 1 (see Figure H).

The ablation results clearly demonstrate that both the proposed variable ordering strategy and the new pruning rules significantly contribute to the performance of FASTiso. Their impact is particularly pronounced on challenging datasets such as SCALEFREE and RAND, where removing either component leads to slowdowns of several orders of magnitude. Importantly, the combination of the two techniques yields substantially larger gains than each component alone, highlighting a strong synergistic effect.

On the Meshes dataset, where FASTiso is not the fastest solver overall, both contributions remain beneficial. The performance degradation observed when reverting to the VF3 ordering confirms that the proposed ordering strategy generalizes well beyond the datasets where FASTiso achieves the best absolute performance. The behavior on PHASE deserves special attention. While the variable ordering strategy of VF3 yields slightly better performance on this dataset, the proposed pruning rules remain effective and do not constitute the limiting factor in this setting.

#### H. Results on MIVIA LDG dataset

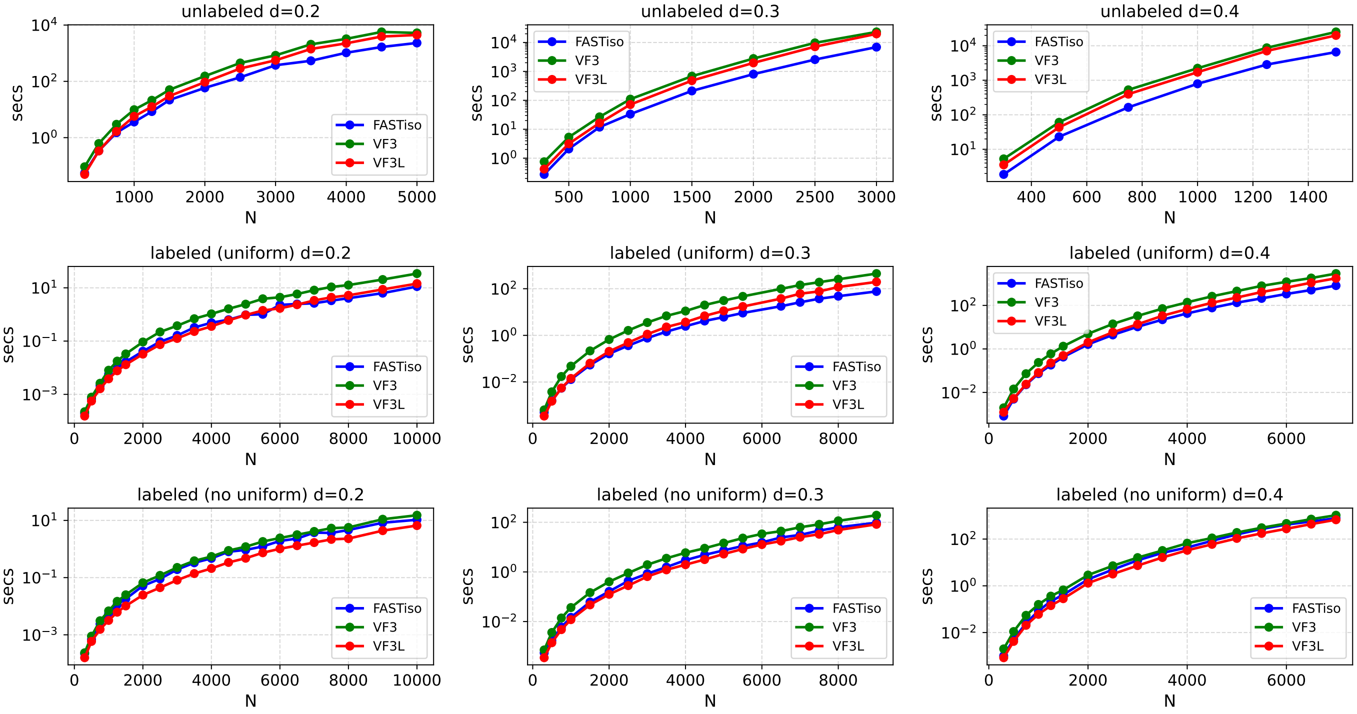


Fig. I. The average matching times for finding all solutions on the graphs from the NIVIA LDG dataset. $d$ represents the density of the graphs, and $N$ is the number of nodes in the target graph.

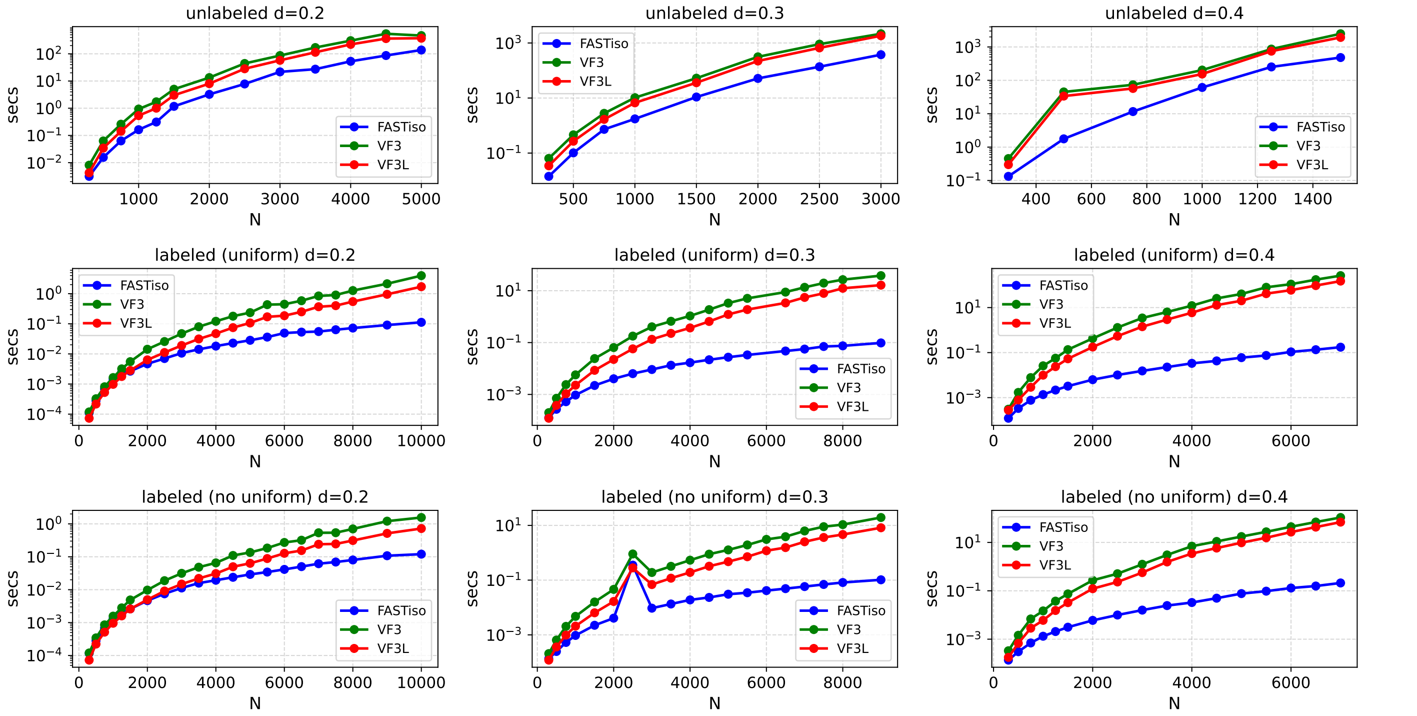


Fig. J. The average matching times for finding one solution on the graphs from the NIVIA LDG dataset. $d$ represents the density of the graphs, and $N$ is the number of nodes in the target graph.

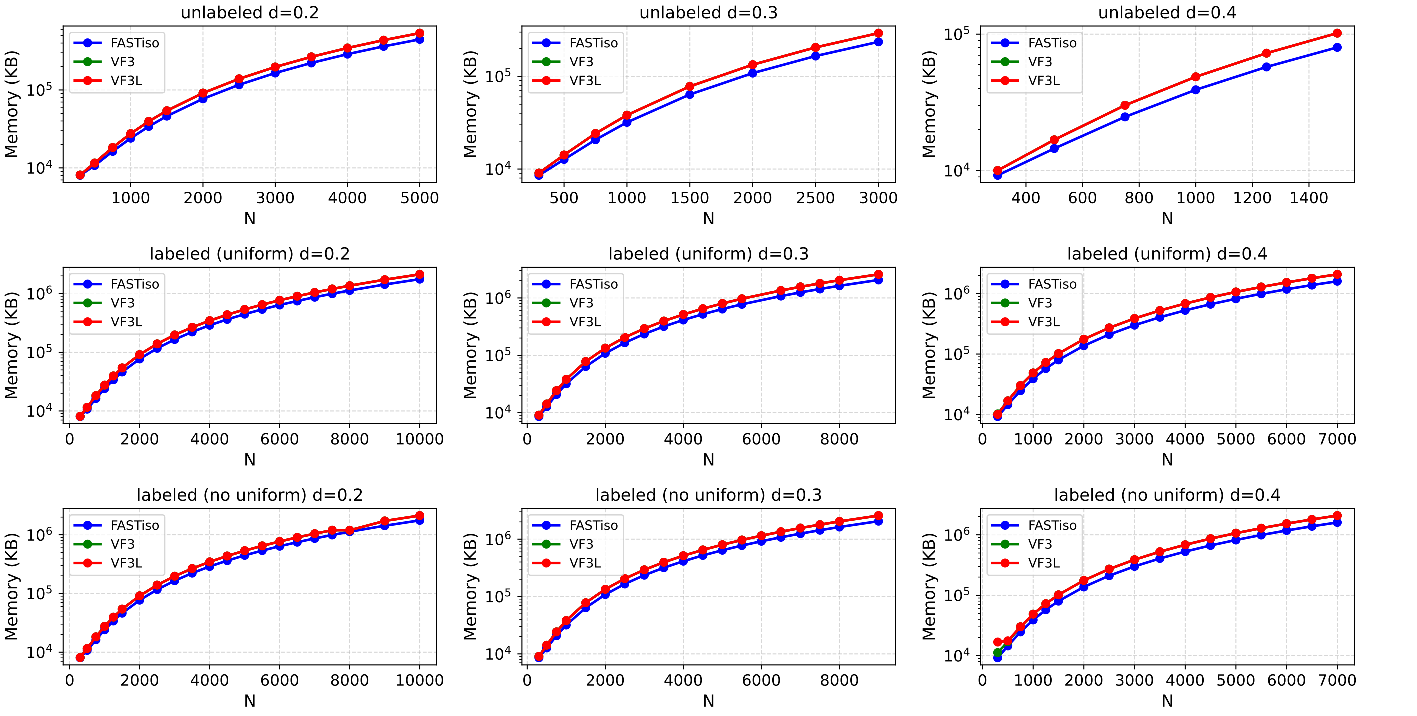


Fig. K. The average memory usage on the graphs from the NIVIA LDG dataset. $d$ represents the density of the graphs, and $N$ is the number of nodes in the target graph.

#### I. Results on Classic benchmark

TABLE 2
Comparison on the Scalefree dataset. Time is in second, #Mem (Mean memory usage in KB).

| solver | Min | 25% | 50% | 75% | Max | Mean | Sum | #Mem |
|---|---|---|---|---|---|---|---|---|
| FASTiso | 1.00E-06 | 8.90E-05 | 2.68E-04 | 5.32E-04 | 6.67E-03 | 4.01E-04 | 4.01E-02 | 7274.70 |
| Glasgow | 1.00E-06 | 1.00E-06 | 2.00E-03 | 6.25E-03 | 2.10E-02 | 5.89E-03 | 5.89E-01 | 10790.96 |
| PathLad+ | 2.78E-03 | 1.71E-02 | 2.64E-02 | 5.68E-02 | 1.89E+00 | 6.72E-02 | 6.72E+00 | 72369.35 |
| VF3 | 3.04E-04 | 1.25E-03 | 3.39E-03 | 5.30E-02 | 7.68E+03 | 1.61E+02 | 1.61E+04 | 7508.00 |
| VF3L | 1.88E-04 | 9.75E-04 | 4.40E-03 | 9.29E-02 | 3.83E+05 | 7.64E+03 | 7.64E+05 | 7456.60 |
| RI | 1.00E-04 | 4.39E-04 | 1.91E-03 | 1.29E-00 | 6.45E+05 | 1.27E+04 | 1.27E+06 | 6761.16 |

TABLE 3
Comparison on the Rand dataset. Time is in second, #Mem (Mean memory usage in KB).

| Target size | Solver | Min | 25% | 50% | 75% | Max | Mean | Sum | #Mem |
|---|---|---|---|---|---|---|---|---|---|
| 200 | FASTiso | 7.10E-05 | 9.25E-04 | 3.63E-03 | 1.34E-02 | 0.37 | 2.91E-02 | 1.74 | 6689.66 |
| | Glasgow | 1.00E-06 | 7.50E-04 | 5.00E-03 | 9.52E-02 | 0.95 | 8.90E-02 | 5.34 | 8530.93 |
| | RI | 6.00E-05 | 2.49E-03 | 3.01E-02 | 0.34 | 2.81 | 0.35 | 21.50 | 6370.00 |
| | VF3L | 7.60E-05 | 6.68E-03 | 7.40E-02 | 0.42 | 2.73 | 0.41 | 24.95 | 6712.40 |
| | VF3 | 1.32E-04 | 3.58E-03 | 4.53E-02 | 0.35 | 4.43 | 0.48 | 29.24 | 6720.40 |
| | PathLad+ | 4.42E-03 | 2.68E-02 | 2.48E-01 | 2.78 | 116.39 | 12.78 | 767.04 | 58203.73 |
| 400 | FASTiso | 5.33E-04 | 9.47E-03 | 0.12 | 1.01 | 358.43 | 8.11 | 730.75 | 7822.17 |
| | Glasgow | 1.00E-06 | 6.35E-02 | 0.33 | 6.56 | 687.01 | 14.30 | 1287.08 | 11709.06 |
| | RI | 5.60E-04 | 0.16 | 1.43 | 37.46 | 531.79 | 36.37 | 3273.38 | 7052.00 |
| | VF3L | 5.55E-04 | 0.22 | 1.96 | 33.86 | 515.91 | 40.61 | 3655.10 | 8147.11 |
| | VF3 | 6.77E-04 | 0.21 | 2.05 | 51.05 | 1105.34 | 66.60 | 5994.84 | 8155.06 |
| | PathLad+ | 1.55E-02 | 1.59 | 125.28 | 225.34 | 7368.86 | 237.27 | 21354.35 | 63736.56 |
| 600 | FASTiso | 3.75E-04 | 1.93E-02 | 0.23 | 4.025 | 578.34 | 16.22 | 1460.35 | 9717.86 |
| | Glasgow | 1.00E-03 | 1.97E-02 | 2.70 | 45.95 | 655.18 | 39.74 | 3577.11 | 18617.91 |
| | RI | 1.16E-03 | 0.17 | 5.80 | 170.78 | 6815.49 | 268.15 | 24134.08 | 8181.37 |
| | PathLad+ | 1.97E-02 | 5.490 | 186.30 | 497.45 | 9359.05 | 569.15 | 51224.07 | 79118.84 |
| | VF3L | 7.69E-04 | 1.09 | 8.12 | 226.38 | 740589.00 | 8528.66 | 767579.51 | 10565.51 |
| | VF3 | 9.76E-04 | 0.85 | 9.62 | 314.23 | 800851.00 | 9326.36 | 839372.58 | 10573.22 |

TABLE 4
Comparison on the BVG dataset. Time is in second, #Mem (Mean memory usage in KB).
Out of 540 instances, FASTiso solved 511 instances, RI solved 510, Glasgow solved 508, VF3/VF3L solved 507, and PathLad+ solved 486

| solver | Min | 25% | 50% | 75% | Max | Mean | Sum | #Mem |
|---|---|---|---|---|---|---|---|---|
| FASTiso | 8.16E-04 | 4.68E-03 | 1.53E-02 | 8.40E-02 | 1.61E+04 | 9.30E+01 | 4.71E+04 | 6973.50 |
| RI | 1.19E-04 | 1.34E-03 | 3.10E-02 | 1.57E-01 | 1.80E+04 | 1.83E+02 | 9.29E+04 | 6659.71 |
| VF3L | 7.80E-05 | 1.62E-03 | 3.57E-02 | 1.88E-01 | 1.80E+04 | 2.34E+02 | 1.19E+05 | 6964.90 |
| Glasgow | 2.00E-03 | 2.75E-02 | 2.17E-01 | 1.91E+00 | 1.80E+04 | 2.45E+02 | 1.24E+05 | 8716.62 |
| VF3 | 1.09E-04 | 2.18E-03 | 4.40E-02 | 2.40E-01 | 1.80E+04 | 3.07E+02 | 1.56E+05 | 6986.09 |
| PathLad+ | 8.71E-04 | 3.96E-02 | 6.66E-01 | 7.50E+01 | 1.80E+04 | 9.22E+02 | 4.68E+05 | 62963.29 |

TABLE 5
Comparison on the M4DR dataset. Time is in second, #Mem (Mean memory usage in KB).
Out of 270 instances, FASTiso and VF3L solved 245, VF3 and RI solved 244, Glasgow solved 243, and PathLad+ solved 207

| solver | Min | 25% | 50% | 75% | Max | Mean | Sum | #Mem |
|---|---|---|---|---|---|---|---|---|
| FASTiso | 1.01E-03 | 2.22E-03 | 6.49E-03 | 6.83E-02 | 1.43E+04 | 1.24E+02 | 3.04E+04 | 7223.41 |
| VF3L | 1.18E-04 | 6.53E-04 | 2.29E-03 | 9.19E-02 | 1.67E+04 | 1.32E+02 | 3.24E+04 | 7230.17 |
| RI | 1.25E-04 | 4.69E-04 | 1.63E-03 | 6.18E-02 | 1.80E+04 | 1.52E+02 | 3.73E+04 | 6780.26 |
| VF3 | 1.82E-04 | 1.00E-03 | 3.57E-03 | 1.37E-01 | 1.80E+04 | 2.38E+02 | 5.83E+04 | 7279.60 |
| Glasgow | 3.00E-03 | 1.20E-02 | 6.70E-02 | 4.47E-01 | 1.80E+04 | 5.20E+02 | 1.27E+05 | 9408.31 |
| PathLad+ | 8.71E-04 | 1.00E-02 | 1.05E-01 | 5.71E+00 | 1.80E+04 | 2.82E+03 | 6.92E+05 | 68911.63 |

TABLE 6
Comparison on the M4D dataset. Time is in second, #Mem (Mean memory usage in KB).
Out of 90 instances, FASTiso, VF3L, VF3, RI, and Glasgow solved 64, and PathLad+ solved 67

| solver | Min | 25% | 50% | 75% | Max | Mean | Sum | #Mem |
|---|---|---|---|---|---|---|---|---|
| PathLad+ | 6.84E-03 | 3.38E-02 | 9.62E-02 | 6.18E-01 | 1.30E+02 | 5.28E+00 | 3.539E+02 | 65869.84 |
| FASTiso | 1.04E-03 | 3.89E-03 | 9.71E-03 | 2.75E-02 | 1.80E+04 | 8.08E+02 | 5.416E+04 | 7068.12 |
| VF3L | 1.69E-03 | 5.14E-03 | 1.56E-02 | 1.01E-01 | 1.80E+04 | 8.09E+02 | 5.417E+04 | 7037.56 |
| RI | 1.06E-03 | 5.20E-03 | 1.10E-02 | 7.26E-02 | 1.80E+04 | 8.10E+02 | 5.427E+04 | 6697.87 |
| VF3 | 1.43E-03 | 6.19E-03 | 1.90E-02 | 8.18E-02 | 1.80E+04 | 8.12E+02 | 5.437E+04 | 7079.93 |
| Glasgow | 3.00E-03 | 1.90E-02 | 6.30E-02 | 2.87E-01 | 1.80E+04 | 8.12E+02 | 5.441E+04 | 9070.25 |

TABLE 7
Comparison on the images-PR15 dataset. Time is in second, #Mem (Mean memory usage in KB)

| solver | Min | 25% | 50% | 75% | Max | Mean | Sum | #Mem |
|---|---|---|---|---|---|---|---|---|
| FASTiso | 8.50E-05 | 4.20E-04 | 1.28E-03 | 2.65E-03 | 3.12E-02 | 2.80E-03 | 6.73E-02 | 8204.50 |
| VF3L | 1.05E-04 | 6.42E-04 | 2.46E-03 | 4.11E-03 | 4.29E-02 | 4.77E-03 | 1.14E-01 | 8817.00 |
| RI | 1.25E-04 | 4.61E-04 | 3.57E-03 | 5.60E-03 | 4.72E-02 | 5.48E-03 | 1.31E-01 | 7431.33 |
| VF3 | 1.20E-04 | 7.15E-04 | 2.71E-03 | 5.39E-03 | 5.66E-02 | 5.79E-03 | 1.38E-01 | 8941.00 |
| Glasgow | 1.77E-04 | 9.57E-04 | 6.24E-03 | 2.50E-02 | 4.51E-01 | 3.07E-02 | 7.37E-01 | 24638.50 |
| PathLad+ | 1.81E-02 | 8.74E-02 | 2.34E-01 | 3.77E+00 | 8.77E+01 | 8.29E+00 | 1.99E+02 | 267093.00 |

TABLE 8
Comparison on the images-CVIU11 dataset. Time is in seconds, #Mem (Mean memory usage in KB)

| solver | Min | 25% | 50% | 75% | Max | Mean | Sum | #Mem |
|---|---|---|---|---|---|---|---|---|
| FASTiso | 1.00E-06 | 1.16E-04 | 1.78E-04 | 3.95E-04 | 8.75E-03 | 5.06E-04 | 3.18E+00 | 7781.12 |
| Glasgow | 3.00E-06 | 8.40E-05 | 2.67E-04 | 1.16E-03 | 1.99E-02 | 1.25E-03 | 6.86E+00 | 19461.49 |
| VF3L | 3.50E-05 | 2.29E-04 | 4.43E-04 | 1.20E-03 | 2.08E-02 | 1.50E-03 | 9.39E+00 | 8278.07 |
| VF3 | 5.60E-05 | 2.65E-04 | 4.52E-04 | 1.19E-03 | 2.63E-02 | 1.50E-03 | 9.41E+00 | 8313.48 |
| RI | 7.00E-05 | 3.61E-04 | 6.30E-04 | 2.08E-03 | 2.46E-02 | 2.19E-03 | 1.37E+01 | 7155.48 |
| PathLad+ | 2.03E-03 | 3.75E-02 | 7.42E-02 | 1.38E-01 | 6.01E-01 | 9.97E-02 | 6.26E+02 | 201004.68 |

TABLE 9
Comparison on the Meshes dataset. Time is in second, #Mem (Mean memory usage in KB). Glasgow and PathLad+ solved all instances, FASTiso failed to solve 1 instance, VF3 failed on 44 instances, RI on 246 instances, and VF3L on 255 instances.

| solver | Min | 25% | 50% | 75% | Max | Mean | Sum | #Mem |
|---|---|---|---|---|---|---|---|---|
| PathLad+ | 1.00E-06 | 4.92E-03 | 1.18E-02 | 5.41E-02 | 9.19E-01 | 6.02E-02 | 1.81E+02 | 114271.95 |
| Glasgow | 1.00E-06 | 1.00E-03 | 4.00E-03 | 2.40E-02 | 2.42E+00 | 9.37E-02 | 2.83E+02 | 14863.61 |
| FASTiso | 1.00E-06 | 1.00E-06 | 1.00E-06 | 1.36E-04 | 1.80E+04 | 5.96E+00 | 1.800077E+04 | 7068.40 |
| VF3 | 4.00E-06 | 6.55E-04 | 1.48E-03 | 3.92E-03 | 1.80E+04 | 3.54E+02 | 1.06E+06 | 7254.85 |
| RI | 1.00E-06 | 2.90E-04 | 7.41E-04 | 1.72E-02 | 1.80E+04 | 1.51E+03 | 4.58E+06 | 7058.94 |
| VF3L | 4.00E-06 | 8.07E-04 | 2.33E-03 | 2.02E-02 | 1.80E+04 | 1.56E+03 | 4.71E+06 | 7254.85 |

TABLE 10
Comparison on the phase dataset. Time is in second, #Mem (Mean memory usage in KB).

| solver | Min | 25% | 50% | 75% | Max | Mean | Sum | #Mem |
|---|---|---|---|---|---|---|---|---|
| Glasgow | 6.76 | 34.33 | 49.36 | 57.74 | 64.33 | 44.63 | 2231.59 | 9197.20 |
| PathLad+ | 184.21 | 234.66 | 249.23 | 262.14 | 272.90 | 244.20 | 11477.75 | 57158.04 |
| FASTiso | 12.72 | 210.84 | 707.50 | 1334.04 | 7598.06 | 1175.53 | 58776.58 | 7068.40 |
| VF3L | 93.53 | 990.70 | 4352.05 | 7723.15 | 17182.40 | 5129.04 | 256452.06 | 7254.85 |
| VF3 | 124.19 | 1084.07 | 4395.70 | 8576.57 | 15720.80 | 5364.50 | 268224.88 | 7266.00 |
| RI | 104.71 | 1398.17 | 5309.96 | 9017.92 | 14858.60 | 5593.71 | 290873.32 | 6605.33 |

##### J. Experiment on isomorphism

This section presents the results of the graph isomorphism experiment using our version of FASTiso for NetworkX, as well as those of VF2 and VF2++.

We generated several types of graphs using NetworkX, with specific parameters defined for each graph model. The graph types used are as follows:

- Random Erdős-Rényi Graphs: These graphs were generated by connecting each pair of nodes with a fixed probability $p = 0.1$.

- Barabási-Albert Graphs: The Barabási-Albert model generates power-law graphs through a preferential attachment mechanism. In this model, new nodes connect to $m$ existing nodes with a probability proportional to their degree. We have set $m$ to 10.
- Watts-Strogatz Graphs: This model generates small-world graphs, where each node is initially connected to its $k$ nearest neighbors in a ring topology. Each edge is then reconnected to a random node with a probability $P$. For our experiment, we define $k = 10$ and $P = 0.1$.
- Bipartite Graphs: We generated bipartite graphs with the same nodes in each set, connecting pairs of nodes with a probability $p = 0.1$.
- Regular Graphs: We have generated 100-regular graphs.
- Power-Law Cluster Graphs: These graphs are generated using a clustering model based on a power-law distribution, where the degree distribution follows a power-law and a local clustering process is added to the graph. We set $m = 3$, the number of random edges to add for each new node, and $P = 0.1$, the probability of adding a triangle after adding a random edge.
- Path Graphs and Cycle Graphs.

For each model, we generated 20 graphs ranging from 500 to 10,000 nodes, with a seed varying from 2 to 20 in increments of 2. The experiments were conducted by setting a time limit of 3600 seconds for processing, and we measured the execution time required to find all the solutions. If an algorithm fails to solve an instance within the given time limit, we consider the time limit as its execution time.

The results are presented in Figure L, where FASTiso is the best in all cases.

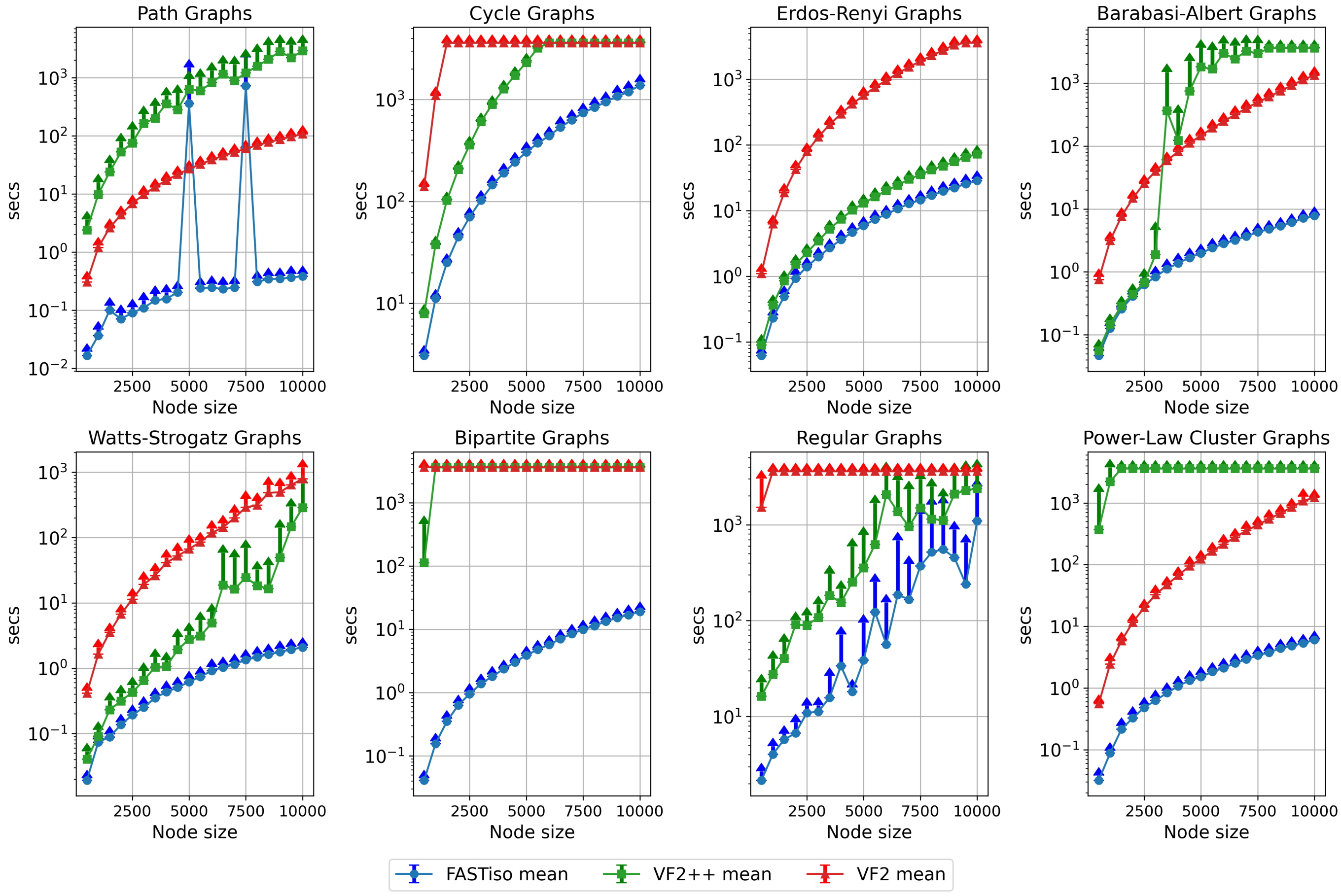


Fig. L. Average matching times for finding all solutions across all datasets
